# Deep learning-driven neuromorphogenesis screenings identify repurposable drugs for mitochondrial disease

**DOI:** 10.1101/2024.07.08.602501

**Authors:** Carmen Menacho, Satoshi Okawa, Iris Álvarez-Merz, Annika Wittich, Mikel Muñoz-Oreja, Pawel Lisowski, Tancredi Massimo Pentimalli, Agnieszka Rybak-Wolf, Gizem Inak, Shiri Zakin, Mathuravani Thevandavakkam, Laura Petersilie, Andrea Zaliani, Barbara Mlody, Annette Seibt, Justin Donnelly, Kasey Woleben, Jose Fernandez-Checa, Diran Herebian, Ertan Mayatepek, Nikolaus Rajewsky, Antonella Spinazzola, Markus Schuelke, Ethan Perlstein, Andrea Rossi, Felix Distelmaier, Ian J. Holt, Ole Pless, Christine R. Rose, Antonio Del Sol, Alessandro Prigione

## Abstract

Mitochondrial disease encompasses untreatable conditions affecting tissues with high energy demands. A severe manifestation of mitochondrial disease is Leigh syndrome (Leigh), which causes defects in basal ganglia and midbrain regions, psychomotor regression, lactic acidosis, and early death. We previously generated isogenic pairs of Leigh cerebral organoids and uncovered defects in neuromorphogenesis. Here, we leveraged on this disease feature to devise drug discovery pipelines. We developed a deep learning algorithm tailored for cell type-specific drug repurposing to identify drugs capable of promoting neuronal commitment. In parallel, we performed a survival drug screen in yeast and validated the repurposable hits on branching capacity in Leigh neurons. The two approaches independently highlighted azole compounds, Talarozole and Sertaconazole, both of which lowered lactate release and improved neurogenesis and neurite organization in Leigh midbrain organoids. Hence, targeting neuromorphogenesis has led to identify potential new drugs for mitochondrial disease and could prove an effective strategy for further drug discovery.

## Introduction

Mitochondrial disease includes rare conditions affecting organs with high energy needs such as the brain, which requires highly functional mitochondrial oxidative phosphorylation (OXPHOS).^1^ One of the most severe forms of mitochondrial disease in children is Leigh syndrome (Leigh) (OMIM #256000).^2^ Leigh, also known as “infantile subacute necrotizing encephalomyelopathy”,^3^ causes lesions in the basal ganglia and midbrain, leading to motor impairment, intellectual disability, and lactic acidosis,^4^ with early death following episodes of metabolic decompensation.^5,6^ The most frequently mutated gene in Leigh is *SURF1* (surfeit locus protein 1, NM_003172.2), an assembly factor for complex IV of the mitochondrial respiratory chain.^7,8^ There is a paucity of effective animal models of Leigh^9^, especially of SURF1 defects, as *Surf1* knock-out (KO) mice failed to develop Leigh-like phenotypes and instead exhibited prolonged lifespan.^10,11^ For this reason, it is currently challenging to identify potential treatments for Leigh.^12,13^

The prevalent view of the pathomechanisms of Leigh is that the disease develops as early-onset neurodegeneration, where neurons die due to excessive production of free radicals because of ineffective OXPHOS.^14^ However, antioxidant therapies failed to show clinical improvements in patients,^12,13^ possibly because antioxidants blunt physiological compensatory responses.^15^ Accordingly, we found no redox imbalance in isogenic induced pluripotent stem cells (iPSCs) model of Leigh carrying SURF1 defects.^16^ Instead, we demonstrated an impairment in neuronal morphogenesis.^16^ Hence, Leigh mutations may interfere with the ability of neurons to form appropriate connections, leading to faulty wiring and reduced resilience to metabolic stress.^17^ Here, we sought to target neuromorphogenesis to devise screening strategies to identify potential candidate compounds for Leigh.

Using single-cell RNA-seq (scRNAseq) datasets of Leigh cerebral organoids (CO), we applied a deep learning (DL) algorithm for cell type-specific drug repurposing and predicted drugs capable of promoting neural commitment in Leigh neural cells. Concurrently, we screened 2,250 FDA-approved drugs for survival in SURF1-KO yeast. The two approaches identified the drugs Sertaconazole and Talarozole, both of which improved neuromorphogenesis in Leigh induced neurons (iNs) and in Leigh midbrain organoids (MO), and ameliorated their growth rate, lactate release, and neuronal metabolism. Collectively, our data indicate that neuromorphogenesis is an effective therapeutic target for Leigh.

## Results

### Neuromorphogenesis screenings in 2D and 3D models of Leigh

We previously demonstrated that Leigh mutations impact the ability of neural progenitor cells (NPCs) to switch to OXPHOS and commit to neuronal fate.^16^ Here, we developed 2D and 3D pipelines to address NPC commitment defects. NPCs were either converted into iNs by overexpressing the transcription factor Neurogenin 2 (NGN2)^18^, or instructed to generate MO^19^ **(Figure 1A)**. Using our high-content analysis (HCA) pipeline to quantify neuromorphogenesis,^20^ we identified significant defects occurring five days after NGN2 induction in Leigh iNs compared to isogenic control (CTL) iNs **(Figure 1B; Figure S1A-B)**. We used scRNAseq to benchmark our MO with respect to previously published MO derived from NPCs^21^ **(Figure S1C)**. We found that our MO contained a high number of tyrosine hydroxylase (TH)-positive midbrain dopaminergic neurons (mDANs) (31 %) already at day 35 **(Figure 1C-D; Figure S1D)**, making this early time point suitable for our analyses. When compared to CTL MO, Leigh MO exhibited aberrant size growth **(Figure 1E)**, increased release of lactate in the medium **(Figure 1F)**, and altered spontaneous calcium activity and metabolic stress-induced calcium signals **(Figure 1G; Figure S1E-L)**. Morphological evaluation of axons (SMI312-positive) and dendrites (MAP2-positive) within Leigh MO confirmed aberrant neurite organization **(Figure 1H)** coupled with reduction of TH-positive neurons **(Figure 1I-J)**.

**Figure 1.**
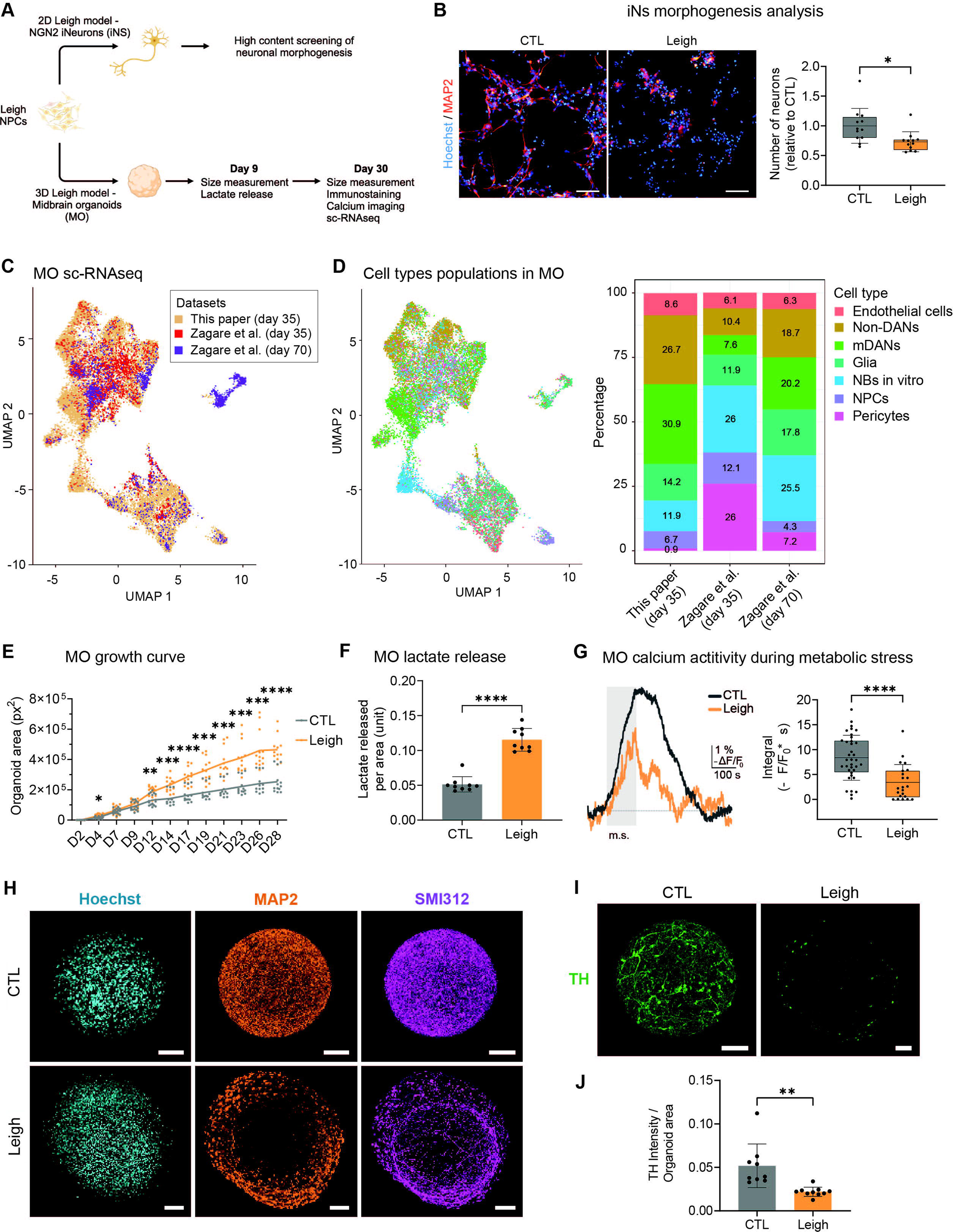
Derivation of 2D and 3D pipelines for targeting neuromorphogenesis in Leigh. **(A)** Schematic view of the 2D and 3D pipelines to assess the ability of NPCs to acquire neuronal fate. **(B)** Outgrowth capacity in induced neurons (iNs) from Leigh and isogenic control (CTL) based on high-content analysis (HCA) quantification. Mean + SD with dots representing 12 biological replicates over 2 independent experiments; *p<0.05, unpaired two-tailed t test. Scale bars: 100 µm. **(C-D)** Unsupervised clustering and population distribution of scRNAseq of our midbrain organoids (MO) (“This paper”) compared to previously derived MO (“Zagare et al”). **(E)** Leigh MO growth rate. Dots represent individual MO over 3 independent experiments; *p<0.05, **p<0.01, ***p<0.005, ****p<0.001, two-way ANOVA. **(F)** Lactate release from MO. Dots represent individual MO over 3 independent experiments. ****p<0.001, Mann-Whitney U test. **(G)** Calcium response in MO following the application of metabolic stress (m.s.) for 2 minutes. Dots represent individual cells (7 CTL MO and 6 Leigh MO) over 2 independent experiments; ****p<0.001, Mann-Whitney U test. **(H-J)** Organization of dendrites (MAP2-positive) and axons (SMI312-positive) and amount of tyrosine hydroxylase (TH)-positive cells in MO. Dots represent individual MO over 3 independent experiments; **p<0.01, unpaired two-tailed t test. Scale bars: 100 µm.

We sought to establish whether aberrant neuromorphogenesis could represent an interventional target in Leigh. We reanalyzed our published scRNAseq datasets of isogenic Leigh CO^16^ to determine key affected populations. Uncommitted precursors showing markers of radial glia (RG) identity were present in both Leigh and CTL CO, while more committed intermediate progenitor cells (IPCs) and neurons were mostly absent in Leigh CO **(Figure 2A-B; Figure S2A-B)**. We therefore reasoned that to counteract this defective neuromorphogenesis, one could forcibly promote neuronal fate in Leigh cells. Pseudotime analysis confirmed that neuronal differentiation was following this directionality RG > IPCs > neurons **(Figure 2C-D)**. We identified differentially expressed transcription factors (DETF) between RG and IPCs and between RG and Neurons **(Figure 2E-F).** We used this defined set of DETFs and a cell type-specific drug perturbation transcriptomics database to develop a DL strategy to uncover drugs capable of triggering cellular conversion from RG directly to neurons (DETF1) or from RG to IPCs and then to neurons (DETF2) **(Figure 2E-F; Figure S2C)**. This strategy highlighted 11 DL-predicted drugs (DLDs) for DETF1 and 17 DLDs for DETF2. Among these 28 DLDs, we selected 9 based on their available safety profile in humans. In parallel, we carried out a rescue screen in Leigh yeast cultures using a library of 2,250 repurposable drugs. The rescue assay was based on survival of wild-type (WT) and SURF1 homologue SHY1^22^ knock-out (ΔSHY) yeast strains upon nutrient removal **(Figure S2F-H)**. Among the top 50 yeast screening drugs (YSDs) **(Figure S2I,** green dots), we selected 4 YSDs belonging to the class of azole-containing compounds, which was the most strongly affected class among rescuer drugs in Leigh yeast **(Figure S2I,** purple dots). We validated DLD1-9 and YSD1-4 in Leigh iNs to assess their impact on neuromorphogenesis **(Figure 2G-H**; **Figure 2J)**. We identified DLD8 (Talarozole) and YSD4 (Sertaconazole) as being capable of increasing neuronal number and neurite length in Leigh iNs in a dose-dependent manner **(Figure 2H-K; Figure S2E; Figure S2K)**. Since the highest concentrations (50 µM) of Talarozole and Sertaconazole showed toxic effects **(Figure S2D; Figure S2J)**, we focused on lower concentrations for downstream analyses.

**Figure 2.**
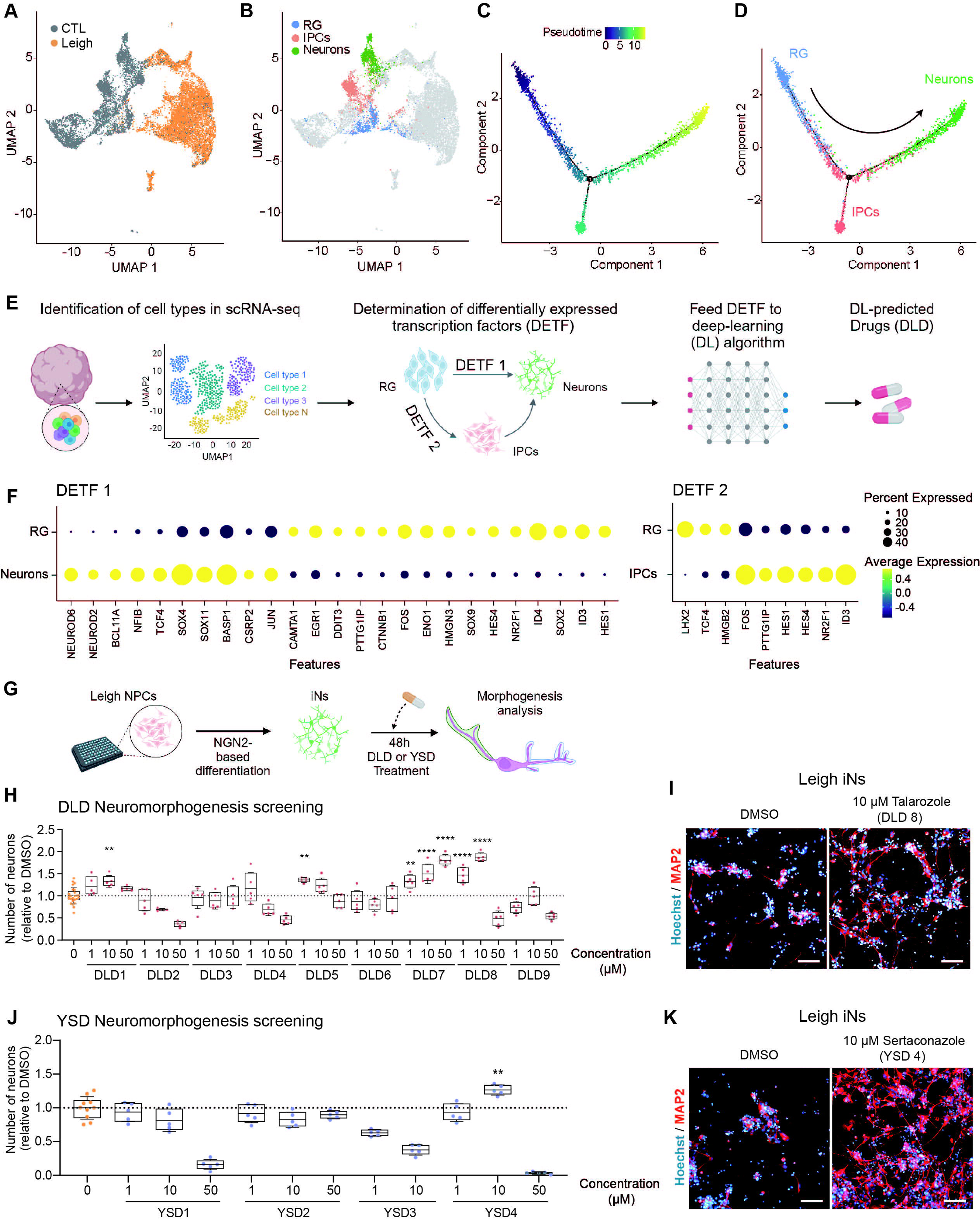
Neuromorphogenesis screenings in Leigh models identified Talarozole and Sertaconazole. **(A-B)** UMAP plots of scRNAseq of cerebral organoids (CO) from Leigh and isogenic control (CTL) showing uncommitted progenitors with radial glia (RG) identity and neural committed cells with identity of intermediate progenitor cells (IPCs) and mature neurons (Neurons). **(C-D)** Pseudotime analysis of the three populations (RG, IPCs, and Neurons). **(E-F)** Schematic of the developed deep learning (DL) algorithm to identify drugs capable of promoting differentially expressed transcription factors (DETF)-specific cell fate conversion. **(G)** Schematic of the 2D neuromorphogenesis screening in Leigh iNs using DL-predicted drugs (DLDs) and yeast screen drugs (YSDs). **(H-K)** DLDs and YSDs impact on neurite outgrowth and representative images for DLD8 (Talarozole) and YSD4 (Sertaconazole). Mean + SD of 5 biological replicates (dots); Statistics is shown only for ameliorating compounds, **p<0.01, ***p<0.005, ****p<0.001, unpaired two-tailed t test, compound-treated Leigh iNs vs DMSO-treated Leigh iNs. Scale bars: 100 µm.

### Identified drugs Sertaconazole and Talarozole rescue Leigh phenotypes

Next, we aimed to determine the effect of the two identified drugs in Leigh MO **(Figure 3A)**. 0.1 µM Sertaconazole or 1 µM Talarozole represented the highest concentrations that could be administrated chronically to Leigh MO without loss of viability **(Figure S3A-B)**. At these concentrations, the drugs rescued the aberrant growth rate of Leigh MO **(Figure 3B)**, reduced their lactate release **(Figure 3C),** promoted the development of TH-positive neurons within Leigh MO, with a significant effect seen for Talarozole **(Figure 3D),** and normalized the disrupted neurite organization of dendrites (MAP2-positive) and axons (SMI312-positive) **(Figure 3E)**.

**Figure 3.**
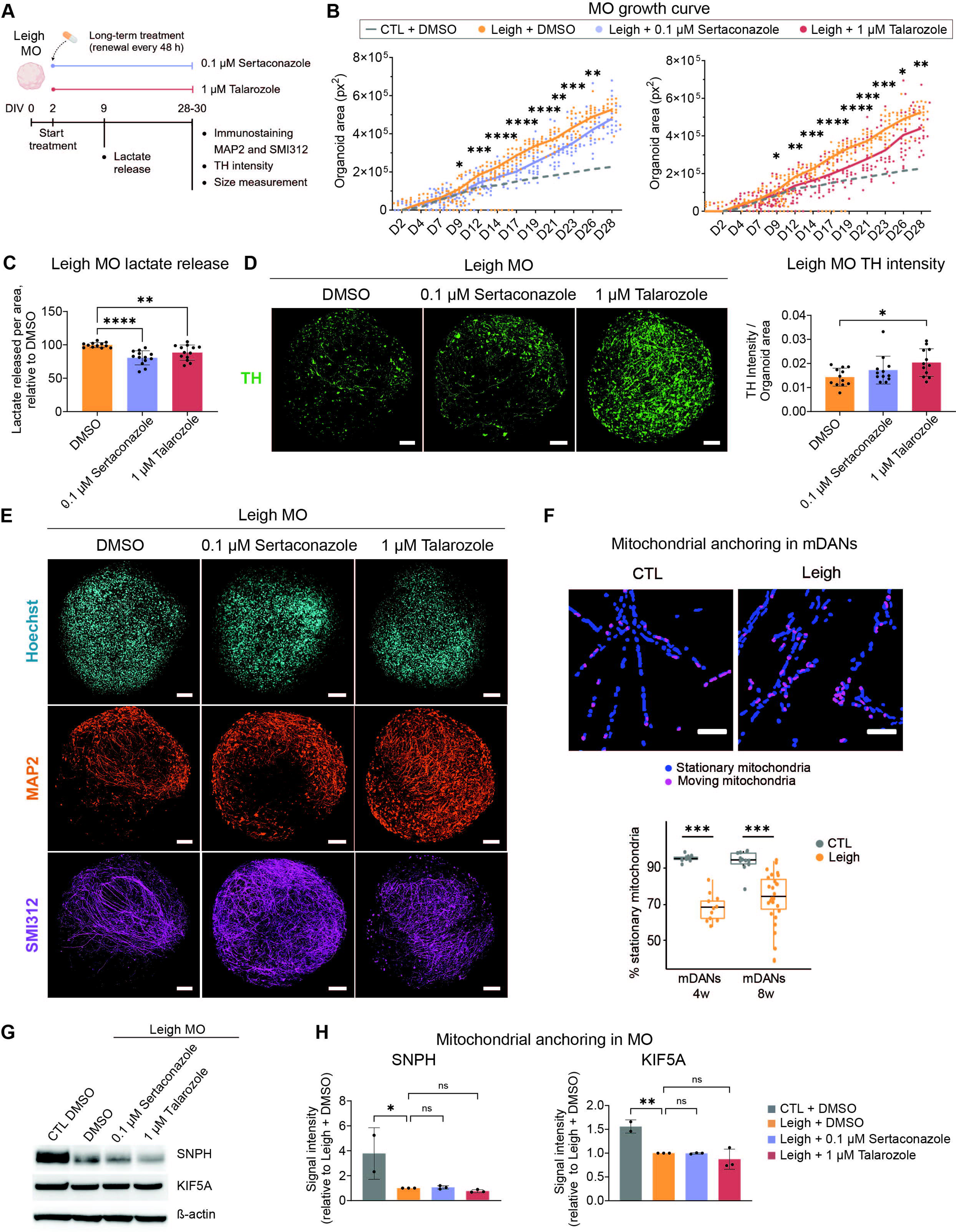
Sertaconazole and Talarozole improve neuromorphogenesis in Leigh organoids without impacting mitochondrial anchoring. **(A)** Schematic of treatment in Leigh midbrain organoids (MO). **(B)** Average growth rate profile of isogenic control (CTL) MO is shown with dotted gray lines. Dots represent individual MO over 3 independent experiments; *p<0.05, **p<0.01, ***p<0.005, ****p<0.001, two-way ANOVA, compound-treated Leigh MO vs DMSO-treated Leigh MO. **(C)** Lactate released in treated Leigh MO compared to DMSO-treated Leigh MO. Dots represent individual MO over 3 independent experiments. **p<0.01, ****p<0.001, Mann-Whitney U test, compound-treated Leigh MO vs DMSO-treated Leigh MO. **(D-E)** Neurite organization and quantification of tyrosine hydroxylase (TH)-positive neurons in treated Leigh MO. Dots represent individual MO over 3 independent experiments; *p<0.05, unpaired two-tailed t test, compound-treated Leigh MO vs DMSO-treated Leigh MO. Scale bars: 100 µm. **(F)** Mitochondrial motility in midbrain dopaminergic neurons (mDANs) from Leigh and isogenic control (CTL) at 4 weeks (4w) or 8 weeks (8w) of differentiation starting from NPCs. Dots represent individual mDANs over 3 independent experiments; ***p<0.005; unpaired two-tailed t test, Leigh mDANs vs CTL mDANs. Scale bars: 100 µm. **(G-H)** Representative immunoblot and related quantification of mitochondrial anchoring proteins SNPH and KIF5A in CTL MO, Leigh MO, and treated Leigh MO. The experiments were performed in 3 independent experiments (dots).

We reasoned that impaired neuromorphogenesis of Leigh might be linked to aberrant mitochondrial motility, as proper anchoring of mitochondria is crucial for neurite branching.^23,24^ Leigh mDANs showed abnormally increased mitochondrial motility with fewer mitochondria that were anchored or stationary **(Figure 3F)**. The increase in mitochondrial movements could not be recapitulated by chemical disruption of mitochondrial viability, which instead caused a decrease in mitochondrial motility **(Figure S3D)**, suggesting the presence of pathological disruption of mitochondrial anchoring. Indeed, the level of mitochondrial anchoring-related genes was lower in Leigh mDANs and Leigh CO compared to CTL **(Figure S3E)**. Leigh MO also displayed reduced expression of mitochondrial anchoring proteins SNPH and KIF5A **(Figure 3G-H)**. However, treating Leigh MO with Sertaconazole or Talarozole did not rescue the expression of anchoring proteins **(Figure 3G-H)**. Hence, promotion of neuromorphogenesis with Sertaconazole or Talarozole might be beneficial for Leigh neural cells despite the persistence of mutation-related defects.

We then focused on neural metabolism and carried out functional metabolic analyses based on scRNAseq of Leigh CO and CTL CO. This kinetic approach comprises the major cellular metabolic pathways of energy metabolism in neural cells and was previously used from proteomics of CO derived from autism spectrum disorders or Huntington’s disease.^25,26^ The technology has been improved to allow its use from scRNAseq datasets. Metabolic genes overlapped between Leigh CO and CTL CO **(Figure S3F-G)**, indicating that it was possible to directly compare their functional metabolic regulation **(Figure S3G)**. Compared to CTL, Leigh CO showed functional defects in energy metabolism (with reduction in oxygen consumption rate, ATP production, and glucose uptake) and in lipid-related metabolism (with defective uptake of fatty acids and ketone bodies) **(Figure 4A-E; Figure S3H-K)**.

**Figure 4.**
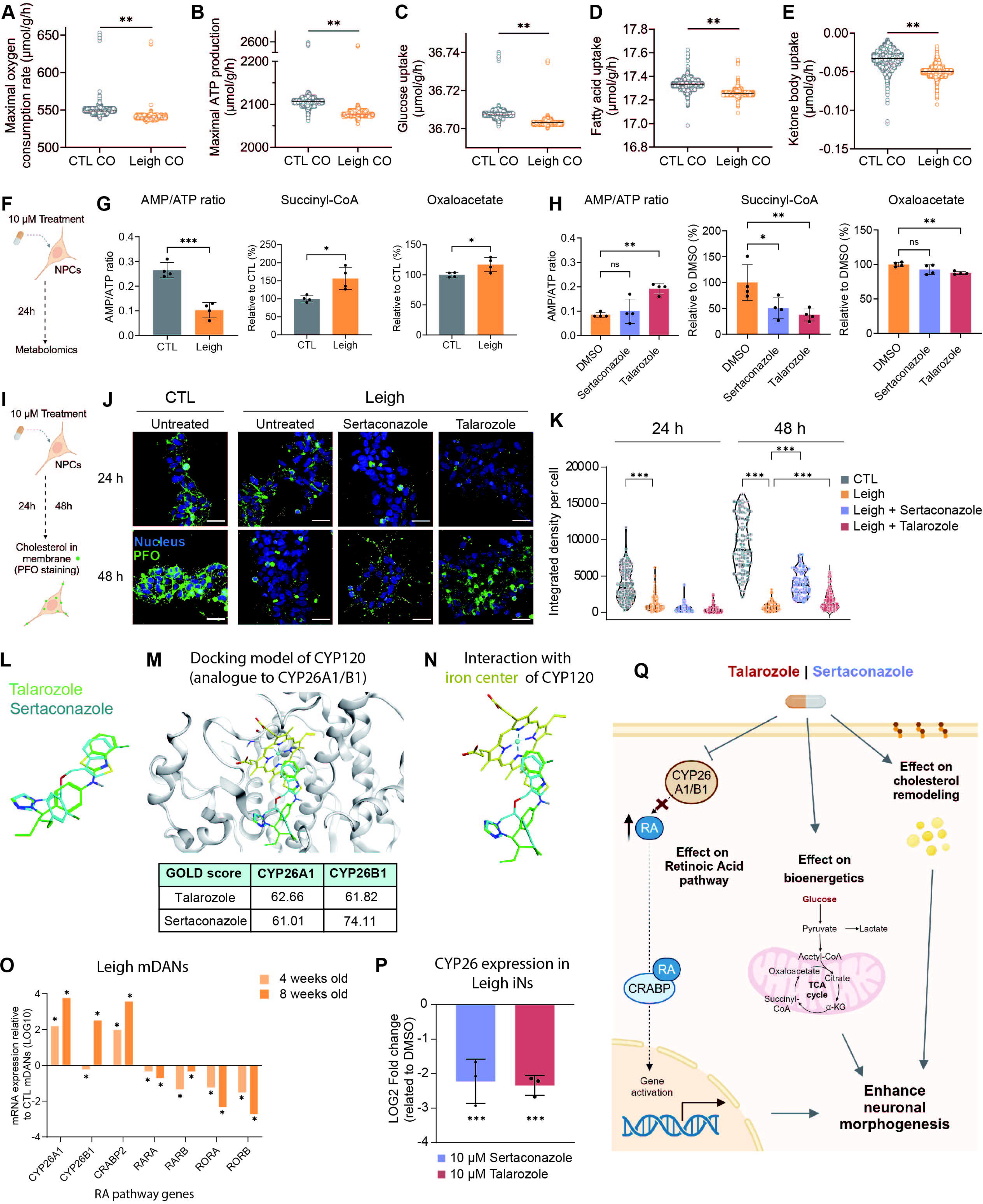
Sertaconazole and Talarozole modulate cholesterol/energy metabolism and retinoic acid (RA) signaling in Leigh models. **(A-E)** Functional metabolic analysis inferred from scRNAseq of cerebral organoids (CO) from Leigh and isogenic control (CTL). Dots represent individual cell values for 3 independent CO scRNAseq samples. **p<0.005; unpaired two-tailed t test. **(F-H)** Targeted metabolomics in NPCs. Dots represent biological replicates over 2 independent experiments; ns: not significant, *p<0.05, **p<0.01, ***p<0.005; unpaired two-tailed t test. **(I-K)** Membrane-bound cholesterol levels measured by PFO staining in CTL NPCs, Leigh NPCs, and treated Leigh NPCs. Dots represent individual values over 3 independent experiments; ***p<0.005; unpaired two-tailed t test. Scale bars: 100 µm. **(L-N)** Predicted docking ability for Sertaconazole and Talarozole to bind the CYP120 pocket. **(O)** Expression of retinoic acid (RA)-related genes in Leigh mDANs compared to CTL mDANs. *p<0.01, fold change expression from bulk RNAseq dataset. **(P)** CYP26 mRNA expression in Leigh induced neurons (iNs) treated with Sertaconazole or Talarozole compared to DMSO-treated Leigh iNs. ***p<0.005; unpaired two-tailed t test. **(Q)** Schematic of proposed mechanisms of action of Sertaconazole and Talarozole in Leigh.

To address the impact of Sertaconazole and Talarozole on energy metabolism, we performed targeted metabolomics in CTL NPCs and Leigh NPCs **(Figure 4F)**. Leigh NPCs exhibited increased glycolysis-related metabolites **(Figure S4B)**, which were not reduced upon treatment with Sertaconazole or Talarozole **(Figure S4D)**. The abnormal levels of tricarboxylic acid cycle (TCA) metabolites such as succinyl-CoA and oxaloacetate **(Figure 4G; Figure S4E)** were instead ameliorated in treated Leigh NPCs **(Figure 4H; Figure S4F)**. The beneficial effects were more evident for Talarozole, which also restored the AMP/ATP ratio of Leigh NPCs **(Figure 4G-H; Figure S4A and S4C)**.

To assess lipid metabolism, we focused on cholesterol, which is essential for neuronal morphogenesis^27,28^ and can be disrupted in mitochondrial disease.^29^ Leigh NPCs exhibited lower membrane-bound cholesterol than CTL NPCs **(Figure 4J-K)**. Treatment with Sertaconazole or Talarozole for 48 hours significantly increased membrane-bound cholesterol in Leigh NPCs **(Figure 4J-K)**. The amelioration was more prominent for Sertaconazole, which is known to modulate the synthesis of ergosterol, the yeast orthologue of cholesterol.^30^ Accordingly, Sertaconazole increased neutral lipid levels in both CTL and Leigh NPCs **(Figure S4G)**.

Lastly, we investigated the molecular structure of Sertaconazole and Talarozole and their *in silico* capacity to bind the main protein target of Talarozole: cytochrome P45026 (CYP26A1/B1) belonging to the RA pathway.^31^ Since the crystal structure of CYP26A1/B1 is not available, we performed the analysis on its homology model CYP120.^32,33^ Sertaconazole and Talarozole showed a biaromatic moiety close to the iron center of heme, with Talarozole positioned in closer proximity to the center than Sertaconazole **(Figure 4L-N, Figure S4I-K)**. Thus, both azoles could bind CYP26A1/B1, although with different affinities **(Figure 4M).** To corroborate this common target, we inspected the expression of RA transcripts in Leigh models. Leigh mDANs exhibited upregulation of enzymes metabolizing RA (e.g. *CYP26A/B1*) and downregulation of RA receptors (e.g. *RORB*) **(Figure 4O)**. Indeed, treating Leigh iNs for 24 hours with either Sertaconazole or Talarozole was sufficient to repress the elevated expression of *CYP26B1* **(Figure 4P)**, indicating that both drugs can modulate this target in Leigh neural cells.

## Discussion

Despite the recent advances in the use of iPSC-based disease modeling and drug discovery, most of the treatments assessed in iPSC models of mitochondrial disease have been compounds suggested by previous studies.^34,35^ This indicates that it is not trivial to identify new therapeutic entities for mitochondrial disease, possibly because of a lack of innovative targets. In Leigh, the culprit for the neuronal pathology is considered to be the death of neurons following oxidative stress.^5^ Antioxidant therapies have however failed to demonstrate clinical efficacy.^13,36,37^ Here, we showed that promoting neuronal generation could represent an effective approach for Leigh. Enabling neuronal growth and proper connectivity might help neurons becoming more resilient against metabolic decompensation.^38^ This could be especially important for dopaminergic neurons, which are strongly affected in Leigh^14^ and are under elevated energy constraints because of their large size and elongated branches.^39-41^

To investigate neuromorphogenesis as a Leigh target, we developed a DL algorithm to identify repurposable drugs promoting neural commitment. Current tools predicting the transcriptional response to chemical perturbations typically rely on datasets based on cancer cell lines.^42-44^ However, cancer cells exhibit signal transduction pathways and metabolic features that are different from those of non-cancer cells^45-47^ and can therefore lead to false predictions for diseases like Leigh which instead affect neuronal cells. We therefore designed a DL-based drug repurposing algorithm that circumvents this problem by using solely datasets from non-cancer cells.^48^ Moreover, our algorithm predicts first genes and then drugs in a cell-type specific manner, thereby allowing to identify also drugs that are outside of training (as was the case for Talarozole). Second, we conducted a survival screen in Leigh yeast. Yeast screens have been used before in the context of neurological and mitochondrial diseases,^49,50^ but their predictive power requires validations in more complex models.^51^ By combining DL prediction and yeast screen with our human 2D and 3D pipelines for neuromorphogenesis, we identified potential repurposable drugs for Leigh.

Sertaconazole and Talarozole shared similarities in chemical structure that might enable them to interact similarly to specific targets. Talarozole is a retinoic acid metabolism blocking agent (RAMBA) and inhibits cytochrome P450 CYP26A1/B1 thereby increasing RA levels in the cell.^52^ Through *in silico* prediction, we found that the antifungal agent Sertaconazole^30^ might also interact with CYP26, suggesting a similar modulation of CYP26A1/B1, which could ultimately influence the RA pathway to promote neuromorphogenesis **(Figure 4Q)**. RA signaling is known for regulating differentiation, organogenesis, and neuronal outgrowth and development.^53-58^ Sertaconazole and Talarozole showed additional beneficial effects on neuronal metabolism by ameliorating OXPHOS, which in turn supports neurogenesis,^17,59-61^ and by modulating cholesterol, which could lead to metabolic reprogramming in NPCs to enhance neuronal generation.^27,28^ We conclude that the effects of Sertaconazole and Talarozole could converge to effect neuronal fate determination in Leigh cells **(Figure 4Q)**. Other drugs belonging to the azole family are under consideration for neurological diseases.^62^ It is thus tempting to speculate that these azole compounds could share similar mechanistic actions on neuronal mitochondria. Indeed, rescuing brain energy metabolism might represent an interventional strategy for common neurodegenerative disorders.^63^

Altogether, we show that neuromorphogenesis represents an effective target process for drug discovery of mitochondrial disease. Furthermore, our DL algorithm might constitute a cost-effective approach to identify drug repurposing candidates in the context of other incurable neurodevelopmental syndromes.

## Limitations of the study

This study was conducted using yeast and iPSC-derived neural models. Additional work is warranted to dissect the systemic *in vivo* impact of these drugs. Sertaconazole and Talarozole are so far only employed topically, even if recent work suggests potential effect for Talarozole and other RAMBAs in reducing inflammation.^64,65^ Since the inflammation modulatory effect of Talarozole is believed to involve the peroxisome proliferator-activated receptor gamma (PPARγ),^66^ it would be important to assess the interplay of Sertaconazole/Talarozole with the PPARγ coactivator-1α (PGC-1α), which we previously found altered in Leigh models.^16^ This work was based only on *SURF1* defects. Additional work is necessary to evaluate whether the two drugs might have positive effects also for other Leigh mutations. Lastly, it remains to be seen whether the combination of Sertaconazole or Talarozole together with other proposed treatments for Leigh^37,67^, including for example the PGC-1α modulator bezafibrate,^16,68^ may potentially lead to synergistic benefits.

## Acknowledgments

We thank Jens Schwamborn (Luxembourg) and his group for providing their published MO datasets. We acknowledge support from the Deutsche Forschungsgemeinschaft (DFG) (PR1527/5-1 and PR1527/6-1 to A.P.; RO5380/1-1 to A.R.; Ro2327/13-2 to C.R.R.; AL2956/1-1 to I.A-M.), the European Joint Programme for Rare Diseases (EJPRD) and Bundesministerium für Bildung und Forschung (BMBF) (AZ. 031L0211 and 01GM2002A to A.P., 01GM2002B to O.P.), the Medical Faculty of Heinrich Heine University (FoKo grant to A.P.), the European Commission’s Horizon Europe Programme (SIMPATHIC #101080249 to A.P.), the United Mitochondrial Disease Foundation (UMDF) and the Leigh Syndrome International Consortium (LSIC) (to A.P), People Against Leigh syndrome (PALS) (to A.P), Foundation Maladies Rare and Association (AMMi) (to A.P), Cure Mito and Cure ATP6 (to A.P.), and MitoHelp (to A.P.). A.Sp. and I.J.H. receive research support from the UK Medical Research Council (MR/X002365/1), Muscular Dystrophy UK (17GRO-PG24-0184-1), and the CHAMP Foundation. A.Sp. is the recipient of a Miriam Marks Senior Fellowship, Brain Research UK (202021-26) and receives funds from the Lily Foundation. I.J.H. is supported by grants from the Instituto de Salud Carlos III PI20/00096 and the Basque Government Department of Health (Osasun Saila, Eusko Jaurlaritzako) (grants 2021111070; 2022333050; 2018111043; 2018222031). M.M.O. was partially supported by the Ikerbasque, Basque Foundation for Science IKUR strategy Neurodegenprot project. We are grateful to Anje Sporbert (Advanced Light Microscopy, MDC, Berlin) for help with the mitochondrial movement recordings, and we acknowledge the Center for Advanced Imaging (CAi) at Heinrich Heine University Düsseldorf for providing access to the PerkinElmer Operetta CLS (DFG grant number INST 208/760-1 FUGG) and Olympus FV3000 microscope.

## Author contributions

Conceptualization, A.P., C.M., A.D.S.; Methodology, C.M., S.O., A.W., M.M-O., L.P., M.T., S.Z., D.H., A.Se., G.I.; Formal Analysis, S.O., T.M.P., C.M., A.Z., I.A-M., B.M., A.R-W.;, J.D.; Resources, E.M., A.P., A.D.S., P.L., A.Sp., E.P., F.D., A.R., N.R., K.W., J.F-C.; Writing – Original Draft, C.M., A.P.; Writing – Review & Editing, A.P., C.M., S.O., A.D.S., O.P., C.R.R., L.P., A.S., I.J.H., I.A-M.; Supervision, A.P., A.D.S., O.P., I.J.H., C.R.R.; Visualization, C.M., A.P.; Funding Acquisition, A.P., A.D.S., O.P.

## Methods

### RESOURCE AVAILABILITY

#### Lead contact

Further information and requests for resources and reagents should be directed to and will be fulfilled by the lead contact, Alessandro Prigione, MD, PhD

#### Materials availability

There are restrictions to the availability of Leigh patient-derived iPSCs used in this study due to the nature of our ethical approval that does not support sharing to third parties without a specific amendment and does not allow to perform genomic studies to respect the European privacy protection law.

#### Data and code availability

The scRNAseq datasets supporting results in this paper are publicly available. The accession numbers are listed in the **Key Resources Table**.

### EXPERIMENTAL MODEL AND STUDY PARTICIPANT DETAILS

#### Induced pluripotent stem cells (iPSCs) and neural progenitor cells (NPCs)

Leigh iPSCs and isogenic control iPSCs were obtained before using Leigh patient-derived cells and CRISPR/Cas9 editing.^16^ Leigh iPSCs carried the *SURF1* mutation c.769G>A (p.G257R) on both alleles in either a Leigh patient background or a healthy control background. The healthy iPSC line was obtained from Dr. Heiko Lickert (Helmholtz Center Munich).^69^ Isogenic control iPSCs carried no *SURF1* mutation in either the patient background or the control background. All iPSC lines were used in accordance with the Ethical Approval obtained by the Medical Faculty of the Heinrich Heine University (Study Number 2020-967_5). iPSC-derived Leigh NPCs and isogenic control NPCs used in this study were generated and characterized before.^16^ All NPCs were cultured as described there, with minor modifications. NPCs grew in matrigel-coated 6-well plates in sm+ medium, consisting of a base media 1:1 ratio of Neurobasal and DMEM-F12 supplemented with N2 (0.5 x), B27 without vitamin A (0.5 x), 2 mM glutamine, Myco-Zap-plus-CL (1 x), 3 µM CHIR 99021, 0.5 μM Purmorphamine, and 150 µM ascorbic acid. Media was exchanged every other day and cells were passaged weekly at a ratio 1:6 to 1:10 using accutase treatment. All NPCs were used in experiments between passage 10 and 25. All cells were routinely monitored to ensure lack of mycoplasma contimation.

### METHOD DETAILS

#### Generation of midbrain organoids (MO)

MO were generated following a published protocol,^19^ with some modifications. To start the generation of MO, NPCs were detached using accutase to obtain a single cell suspension. Cells were then counted using an automatic cell counter, with a gating size of 5-19 μm, and seeded onto low-attachment U-bottom 96-well plates at a density of 9,000 cells/150 μl media per well. Seeding media was composed of a basal media containing 1:1 ratio DMEM-F12:Neurobasal, N2 (0.25 x), B27 without vitamin A (0.25 x), glutamax (1 x), MycoZap-Plus-CL (1 x), 0.4 % (w/v) polyvinyl-alcohol (PVA) supplemented with the following small molecules: 0.5 μM Purmorphamine, 3 μM CHIR 99021 and 100 μM ascorbic acid. After 2 days, seeding media was replaced with a basal media supplemented with 100 µM ascorbic acid, 1 µM Purmorphamine, 1 ng/ml BDNF, and 1 ng/ml GDNF, to induce ventral patterning. Four days later, the media was exchanged to a maturation media, composed of the basal media supplemented with 100 µM ascorbic acid, 2 ng/ml BDNF, 2 ng/ml GDNF, 1 ng/ml TGF-β3, and 100 µM dibutyryl-cAMP. On day 6, a single dose of 5 ng/ml Activin A was added. From day 7 on, maturation media (without Activin A) was refreshed 3 times a week. For experiments with chronic treatment, MO were treated from day 2 until the day of the assay (either until day 9 or day 28-35) with either 0.1 µM Sertaconazole, 1 µM Talarozole or DMSO (vehicle). DMSO samples contained 0.1 % (v/v) DMSO, which is equivalent to the amount of solvent in the samples that received compound treatment. Treatments were refreshed upon every media change every other day.

#### MO growth rate analysis, viability assay, and lactate release

To measure organoid size, pictures were taken every other day with a Nikon Eclipse TS2 inverted routine microscope using a 4 x objective and ensuring a good contrast between the background and the organoid. Organoid area and perimeter were measured using CellProfiler. Color images were first converted into grayscale images using the modules “Color to gray” and “Invert for printing”. Next, grayscale images were used as an input for the module “Identify Primary Objects” to identify a bright object (organoid) of a diameter ranging 300-1200 px on a dark background. The properties of identified objects could then be measured using the module “Measure Object Size Shape”. Undesired identified objects (such as debris) were filtered out using the “Filter Objects” module, based on their compactness, with a maximum value of 4. After filtering, the module “Measure Object Size Shape” was again used to measure the properties of the filtered identified objects (organoid). The results of this module were exported to Excel file for analysis. The organoid area measured at day 2 was used as a reference starting point and therefore subtracted from each individual value obtained. thereafter. At least 5 organoids per cell line for each time point from 3 independent experiments were analyzed. For the viability assay, control MO were treated from day 2 to day 11 every second day with different concentrations (0,1-1-5 and 10 μM) of Talarozole or Sertaconazole. DMSO control samples contained 0.1 % (v/v) DMSO, which is equivalent to the amount of solvent in the samples that received compound treatment. On day 9, organoid viability was measured using CellTiter-Glo cell viability assay following a previously published protocol.^19^ Briefly, the media of each well was adjusted to 55 μl and the plates with MO together with the kit’s reagents were brought to RT for 30 min. Next, an equal volume of 55 μl of Celltiter-Glo reagent was added and samples were shaken on a Thermomixer at 900 rpm for 5 min. Afterwards, plates were incubated for 25 min at room temperature (RT) protected from light. The contents of the plate were transferred to white/opaque 96-well plates with 2 technical replicates (20 μl each) per sample. Luminiscence of each sample was recorded using a Tecan plate reader. Extracellular lactate released by MO in the media was measured by collecting 100 μl of maturation media at day 9 of MO generation. Media from at least 5 individual MO from 3 different batches was collected for analysis and stored at -20 °C until used. The manufacturer’s instruction from the Lactate Assay commercial kit was followed for measuring the absorbance of each sample using a clear 96-well microplate assay and Tecan microplate to read the absorbance at 580 nm. The calculated amount of lactate per well was normalized to the respective organoid size (see *MO growth rate analysis* section) and expressed as fold-change compared to control (when untreated) or to DMSO-treated sample (when treated with Sertaconazole and Talarozole) of their respective batch.

#### Single-cell RNA sequencing (scRNAseq) analyses

35 days-old MO from iPSC control line were dissociated for scRNAseq using the Papain Dissociation System. Papain was prepared by adding 5 mL of EBSS to one papain vial and incubating for 10 min at 37 °C. Additionally, 500 μl of EBSS was added to one DNAse vial and gently mix. Half of the volume of DNAse in EBSS was then added to the papain vial resulting in a final concentration of approximately 20 units/ml of papain and 0.005 % DNase. About 48 individual MO were collected onto a 6 well-plate, washed with PBS, and incubated with 2 mL of pre-warmed papain/DNase solution for 5-10 min at 37 °C in an orbital shaker. Using a 1 mL pipette, the organoids were gently triturated by mixing up and down and placed back into the incubator for another 5-10 min for further enzymatic digestion. The digested organoids were then collected in a 15 mL tube and 5 mL of ovoalbumin was added. Using a 225 mm polished glass pipette, the mixture was further triturated until mostly dissociated. The cell suspension was then filtered through a 30 μm cell strainer and transferred to a new 15 mL tube. The cell concentration was determined, and cells were pelleted by centrifuging at 300 x g for 5 min. After removing the supernatant, the pellet was resuspended in 200 μl of ice-cold PBS and 800 μl of freezing-cold 100 % methanol was added. The cell suspension was snap-frozen in liquid nitrogen and stored at -80 °C until library preparation. After rehydration of the samples, libraries were generated using 10x Chromium and sequenced with Illumina NovaSeq 6000 platform. Samples were subjected to scRNAseq using the 10× Genomics Chromium Single Cell 3’ Gene Expression system with feature barcoding technology for cell multiplexing. Demultiplexed, raw paired-end scRNAseq data was processed using the Cell Ranger (v. 7.10) from the 10x Genomics. Data was analyzed in three biological replicates, where each replicate was composed of 48 individual MO. Sequencing was carried out in two separate batches. In the first batch, replicates 1 and 2 were processed, while replicate 3 was processed in the second batch. In this manner, we assessed reproducibility of the sequencing results across different independent runs. For comparing our MO to other MO generated before, we employed scRNAseq datasets of MO that were previously generated at different time points.^21^ The datasets were obtained from GSE133894. Cell type was annotated following the approach described before^19^ where eight cell types were determined based on the expression levels of their marker genes (Endothelial cells, non-mDANs, mDANs, Glia, Nbs in vitro, NPCs and Pericytes). The percentage of each cell type with respect to the total cell counts was computed. Integration of the two datasets was performed using the *FindIntegrationAnchors()* function described before^65^ with default parameters using the Seurat R package (v. 4.2.0).^70^ scRNAseq data of Leigh cerebral organoids (CO) and isogenic control CO were carried out before.^16^ Datasets were obtained from GSE126360. Pseudotime analysis was performed on the RG, IPCs and Neurons populations in those scRNAseq data using Monocle3 R package.^71^ Input genes were differentially expressed genes among the three populations with the adjusted p value cutoff <= 0.05. The DDRTree algorithm was used for the inference.

#### Immunostaining

Free-floating MO were fixed with 4 % PFA for 20 min at RT, followed by three 10 min washes with PBS. Organoids were blocked and permeabilized with a solution consisting of 3 % BSA, 0.5 % Triton-X-100 and 0.05 % sodium azide in PBS, for 1 h at RT. Primary antibodies were diluted in blocking solution and incubated overnight at 4 °C. Afterwards, they were washed 3 times with PBS and incubated overnight at 4 °C with secondary antibodies at a 1:1000 dilution in blocking solution. Next, organoids were incubated for 1 h at RT with Hoechst 33342 (1:1000) and washed three times with PBS. Images were acquired using the ZEISS Axio Observer microscope with an apotome 3 as a z-stack, which were then deconvoluted using the Zeiss blue software default settings and z-projected with maximum intensity. Antibodies are reported in the **Key Resource Table.** For quantification of tyrosine hydroxylase (TH) within MO, immunostained MO z-projected images were analyzed using Columbus software (v. 2.9.0). Prior to analysis, images were downscaled using the greyscale function in Photoshop (v. 12.1 x64) and resized from 300 to 72 pixels per inch. The images were converted from 8-bit to 16-bit channels to retain image information and saved as .tif files. An image analysis pipeline was established using different building blocks. MO objects were identified based on the TH channel using the “Find Image Region” block. The “Fill Region” and “Border” functions within the “Select Region” block were utilized to cover the entire organoid area. Within this defined area, the intensities of each channel were calculated using the “Calculate Intensity Properties” block and given as the average pixel intensity. Furthermore, morphological parameters such as area, width, length (all measured in µm), and roundness were assessed using the “Calculate Morphology Properties” block. For the comparison of Leigh MO and control MO, we calculated the ratio of intensities of respective channels and the area of the individual MO.

#### Intracellular calcium imaging of MO

33-37 days-old MO were stained with the membrane-permeable form of the calcium indicator Fura-2 (Fura-2 AM; 1 mM stock solution in 20 % pluronic/DMSO; 15 µM final concentration) dissolved in artificial cerebrospinal fluid (ACSF), containing (in mM): 138 NaCl, 2.5 KCl, 2 CaCl_2_, 1 MgCl_2_, 1.25 NaH_2_PO_4_, 18 NaHCO_3_, and 10 glucose (pH 7.4; osmolarity ∼310 mOsm/l) at 37 °C in a humidified atmosphere with 95 % O_2_ and 5 % CO_2_ for 30 min, followed by additional 30 min in ACSF without dye. A single MO was placed onto an Omnipore membrane filter and stabilized by a grid in the recording chamber. MO were continuously superfused (2–2.5 ml/min) with ACSF bubbled with 95 % O_2_5 % CO_2_ and allowed to equilibrate for ∼10 min before starting recordings. Experiments were carried out at 37 ± 1 °C. Wide-field calcium imaging was performed using an upright microscope equipped with a 40 x/N.A. 0.8 water immersion objective. Fura-2 AM was alternately excited at 380 and 400 nm using a monochromator and images were acquired at a frequency of 1 Hz with a complementary metal–oxide–semiconductor (CMOS) camera. Fura-2 AM emission was collected at 510 ± 40 nm from regions of interest (ROIs) manually drawn around cell bodies. Background correction was performed as previously described.^72^ Spontaneous calcium activity was analyzed during a 5 min recording period, in which no other manipulation was performed. Cells with calcium signals ≥ 2x standard deviation (SD) were considered active. Metabolic stress was induced by perfusing preparations with a glucose-free ACSF containing 2 mM 2-deoxyglucose and 5 mM sodium azide for 2 min. Changes in fluorescence were normalized to the baseline and are expressed as percentage change.

#### Deep learning (DL)-based drug prediction

To predict genes that need to be perturbed for converting one cell fate state into another desired state, a feedforward artificial neural network (FNN) model was trained using the datasets retrieved from ChemPert.^48^ We chose ChemPert since it consists of non-cancer perturbation transcriptomics datasets, which were shown to be more suitable for non-cancer cell studies than cancer-based datasets such as CMAP^42^ and LINCS.^43^ The input data for training the FNN model was composed of: 1) differentially expressed transcription factors (TFs) upon cell stimulation by single perturbagens (both chemical and with protein agents), 2) initial (unperturbed) gene expression state of the cell. The latter was included to take into account cell-type specificity, which plays a critical role in determining transcriptional responses. The training output was the known direct target genes of perturbagens, which are also available in ChemPert. In total, we used 82270 transcriptomics datasets in response to 2566 unique perturbagens across 167 non-cancer cell types for the model training. In the next step, the predicted target genes were compared to the known target genes of 5693 perturbagens and the optimal perturbagen combinations were computed by maximizing the number of true positive (predicted) target genes, while minimizing the number of false positive (unpredicted but targeted by perturbagens) target genes. The FNN model performance was optimized by iterative cross validation. In our previous CO scRNAseq data of Leigh CO and isogenic control CO,^16^ we identified cell clusters for SURF1-corrected NPCs and neurons. We then performed differential expression analysis between these two cell clusters and the DETFs. The gene expression data of the NPCs were fed to the FNN model to identify perturbagens that could induce and accelerate the neurogenesis of Leigh NPCs. The model predicted 27 chemical compounds, 9 of which were short-listed for further experimental evaluation based on the prior knowledge of their biological effects.

#### DL algorithm

A feedforward neural network (FNN) model was trained using the perturbation transcriptomics datasets retrieved from ChemPert^48^ . Signaling proteins and their molecular interaction data were obtained from the following sources: protein-protein interactions from iRefIndex, ^73^ signaling networks from OmniPath,^74^ ReactomeFI,^75^ KEGG,^76^ NicheNet,^77^ and ligand-receptor interactions.^77-88^ Transcription factors (TFs) were obtained from AnimalTFDB 3.0 (http://bioinfo.life.hust.edu.cn/AnimalTFDB2/).^89^ The perturbagens from ChemPert were further filtered by removing the ones targeting apoptosis-related proteins and those not targeting any signaling protein. The FNN model training data consisted of: 1) DETFs between initial and end cell states of each perturbation transcriptomics dataset, 2) gene expression of initial cell state. The FNN model output was direct target proteins of perturbagens. The model algorithm consists of four major steps described below.

Step 1: Gene expression value of each signaling protein p_j_ in the initial cell state is transformed by

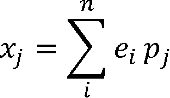

where e_i_ is the gene expression of each direct upstream neighbour protein defined in Step 1. After computing this value for all signaling proteins, the normalization was performed by:

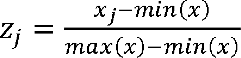

where x is the entire set of transformed signaling protein values. Differentially up- and down-regulated TFs were labeled with values of 1 and -1, respectively. 0 was assigned to all non-DETFs. Perturbagens’ effect on direct protein targets –activation, inhibition or unknown–, were labeled with 1, −1 and 2, respectively.

Step 2: The transformed signaling protein values and DETF values are concatenated into one vector and fed to the FNN model for training. The number of hidden layers and nodes were optimized and set to 3, and 1024, 512, 1024 for hidden layers 1, 2, 3, respectively. Batch normalization is applied after each layer to enhance model stability. Backpropagation is performed using the Adam optimizer with a learning rate set to 0.005 to optimize the model using the cross-entropy loss function. The softmax function is used for the activation function in the output layer. The training epoch number is set to 1000.

Step 3: Pathway enrichment analysis is carried out on predicted target signaling proteins using the Reactome database. Pathways with fewer than 3 or more than 500 signaling proteins are excluded before analysis. Subsequently, enriched pathways with adjusted p-value <= 0.05 and containing more than 3 predicted signaling proteins are retained for the next step.

Step 4: For each enriched pathway, predicted signaling proteins present in that pathway are compared to the direct target proteins of perturbagens to identify optimal perturbagen combinations by iteratively maximizing the Jaccard index:

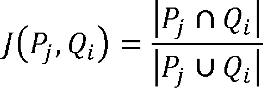

where p_j_ is predicted target proteins in pathway 1 and Q_i_ is target proteins of perturbagen *i*. First, a “seed” perturbagen is matched to the predicted signaling proteins and the Jaccard index is computed. Then the next perturbagen is matched to the remaining untargeted predicted signaling proteins and the Jaccard index is re-computed and if the index is higher than the previous value, then the perturbagen is retained. This process is repeated until no increase in the Jaccard index is observed for all perturbagens. Note, when the effect of predicted proteins is “unknown”, these proteins are considered both activation and inhibition. Perturbagens with the Jaccard index less than 0.1 are discarded.

#### Neuromorphogenesis screenings in iNs

To generate iNs, NGN2 lentiviruses were produced using HEK 293 cells seeded at a 70 % confluence in a 150 cm^2^ dish in DMEM medium.^90^ Once attached, the medium was refreshed with DMEM supplemented with 25 μM Chloroquine and cells were transfected following Lipofectamine 2000 manufactureŕs protocol using the following plasmid mix: 8.1 µg pMD2.G, 12.2 µg pMDLg-pRRE, 5.4 µg pRSV-Rev, 5.4 µg FUW-M2-rtTA, 14.3 µg TetO-FUW-NGN2, 14.3 µg TetO-FUW-EGFP.^91,92^ The next day, the medium was exchanged with DMEM-F12 containing 10 µM sodium butyrate. The supernatant containing viral particles was collected 24 and 48 h later. To concentrate the virus particles, the supernatant was first centrifuged at 500 x g for 10 min to remove cells and debris and then mixed with 1 volume of cold Lenti-X concentrator to every 3 volumes of lentivirus-containing supernatant. This mixture was kept overnight at 4 °C and further centrifuged at 1,500 x g for 45 min at 4 °C. The obtained pellet containing the virus was diluted 1:100 in PBS. Virus aliquots were stored at -80 °C until further use. For neuronal morphogenesis screening, NPCs were seeded onto matrigel-coated 96-well black greiner-bio plates at a density of 40,000 cells/well in 100 μl of sm+ (see *iPSC-derived neural progenitor cells* section for media composition). The next day, the media was refreshed with sm+ supplemented with 4 μg/ml polybrene and 2.25 x 10^6^ transducing units of NGN2 lentivirus. After overnight incubation, NPCs were washed 3 times with PBS and the media was refreshed with sm+ containing 2 μg/ml of doxycycline to start neuronal induction (day 0). On day 3, medium was exchanged to NGN2 medium, consisting of Neurobasal, B27 supplement with vitamin A (0.5 x), NEAA (1 x), 2 mM Glutamine, MycoZap-Plus-CL (1 x), 10 ng/ml human NT-3, 10 ng/ml human BDNF supplemented with 2 μg/ml doxycycline. On day 3, treatments with DLDs and YSDs were administered in the media using 3 different concentrations (1, 10, and 50 μM). Final concentration of DMSO in the media was either 0.1 % or 0.4 % (for 50 µM treatment of DLD4 and DLD5), which was equivalent to the amount of solvent in the samples that received compound treatment. On day 5, after 48 h incubation with the compounds, iNs were fixed by adding equal volumes of 8 % PFA to the medium. Cells were incubated in a final dilution of 4 % PFA for 20 min and washed 3 times with PBS for 10 min/wash. For staining, iNs were blocked and permeabilized for 1 h at RT with blocking solution (3 % BSA, Triton-X-100, 0.05 % sodium azide in PBS). Primary antibody incubation with anti-MAP2 (1:1000) in blocking solution was carried out overnight at 4 °C. Afterwards, they were washed 3 times with PBS and treated with Alexa Fluor Anti-Guinea pig AF568 secondary antibody (1:1000) and Hoechst 33342 (1:2500) in blocking solution for 1 h at RT. Next, after 3 washes with PBS, wells were filled with 0.05 % sodium azide in PBS and stored at 4 °C wrapped with parafilm until imaged. Antibodies are reported in the **Key Resource Table**. iNs were imaged with the high-content microscope Operetta using the 0.4 NA 20X air objective. For each well, 20 areas across the well were selected for imaging. The areas remain constant for each plate analyzed. Images of Hoechst and MAP2 channel were then analyzed with CellProfiler^93^ to analyze neuronal morphogenesis as previously described^18^ Data was summarized by calculating the mean per well. To compare different plates, numbers were compared to the mean of the DMSO-treated wells.

#### Yeast drug screen

Wild-type (WT) and SURF1-KO mutant (ΔSHY) strains were assessed for growth deficiency in a non-fermentable growth medium (YP 2 % lactate) at 30 °C. A 384-well optimization assay was developed: 1) to identify the right starting density of cells that accentuates the growth difference phenotype and best assay time, 2) to establish that the assay has a good Z’, which is a measure of assay quality and the likelihood of false positives and negatives in the screen. In this assay, the Z’ score was 0.7 indicating that the assay quality is excellent. Overnight cultures were diluted to an OD of 0.025. 50 nl of 10 mM test compounds were added to columns 3-22 of 384-well plates, while 50 nl of DMSO was added to control columns 1-2 and 23-24. Mutant and WT cell suspensions were adjusted to an OD of 0.025, and 25 µl of mutant cells were added to columns 1-22, and WT cells to columns 23-24. The plates were spun at 250 x g for 1 min, sealed with a gas-permeable seal, and covered with a water-filled lid, then incubated at 30 °C for 24 h. After incubation, seals were removed and plates were equilibrated at RT for 15 min. 25 µl of Bactiter-glo solution was added to each well, plates were shaken for 2 min, incubated for 10 min, and luminescence was read using an Envision plate reader. A list of hits was generated, with the top 2 % selected for further analysis. Test compounds used in yeast drug screening belong to the UCSF’s Pharmakon library, comprising 1,600 drugs, and an additional 200 unique bioactives. All compounds are FDA-approved with an existing safety record.

#### Functional metabolomics based on CO scRNAseq

For the functional metabolic analysis of Leigh CO compared to isogenic control CO, we employed the Quantitative System Metabolism (QSM) pipeline developed by Doppelganger Biosystem GmbH, as described before.^94^ QSM data analysis used quantitative information on the expression levels of metabolic proteins (enzymes) to determine metabolic profiles, metabolic states and capacities, and metabolic fluxes. The kinetic model includes cellular metabolic pathways related to energy metabolism, electrophysiological processes, and membrane polarization information, including the mitochondrial membrane potential membrane, the utilization of the proton motive force, and the transport of ions across the membranes. Time-variations of small ions were modelled by kinetic equations of the Goldman-Hodgkin-Katz type.^95^ Numerical values for kinetic parameters of the enzymatic rate laws were taken from reported kinetic studies of the isolated enzyme. Maximal enzyme activities (Vmax values) were estimated based on functional characteristics and metabolite concentrations of a healthy tissue. Hormone-dependent regulation of the central energy metabolism by reversible enzyme phosphorylation was taken into account by a phenomenological model of insulin and epinephrine dependent changes of plasma glucose and changes in the phosphorylation state of interconvertible enzymes as described in.^96^ Expression profiles normalized to healthy neuronal tissue were used to scale the maximal activities of enzymes and transporters for each cell. QSM was used to calculate energetic capacity at maximal level at physiological nutrient and oxygen concentrations. The interdependence between plasma glucose, plasma hormone and plasma fatty acid concentration was taken into account by using a sigmoid Hill-type function describing the experimentally determined glucose-insulin and glucose-fatty acid relations.^97^ Energetic capacities were evaluated by computing the changes of metabolic state elicited by an increase of the ATP consumption rate above the resting value.

#### Generation of midbrain dopaminergic neurons (mDANs)

The generation of mDANs from NPCs was performed following a previously published protocol.^16,98^ To start differentiation, NPC media was switched to a mixture of 1:1 of Neurobasal:DMEM-F12, N2 (0.5 x), B27 with vitamin A (0.5 x), 2 mM glutamine and Mycozap-Plus-CL (1 x) supplemented with 200 μM ascorbic acid, 100 ng/ml FGF8 and 0.5 μM Purmorphamine for 7 days. The following two days, the concentration of ascorbic acid and purmorphamine was brought down to 100 μM and 0.25 μM, respectively. On day 9, cells were passaged with accutase and seeded onto matrigel-coated plates with 10 μM ROCK inhibitor in maturation media, made of a mixture 1:1 Neurobasal:DMEM-F12, N2 (0.5 x), B27 with vitamin A (0.5 x), 2 mM glutamine, Mycozap-Plus-CL (1 x), with the addition of 200 μM ascorbic acid, 10 ng/ml BDNF, 10 ng/ml GDNF, 500 μM dibutyryl-cAMP, and 1 ng/ml of TGF-β3. The media was refreshed every other day, and the neurons were kept in culture for 4 and 8 weeks to reach different maturation stages.

#### Mitochondrial movement in mDANs

mDANs cultures were grown on 35 mm dishes with coated bottom and 1.5 coverslips. 25 nM MitoTrackerRed CMXRos was added for 10 min and then replaced with DA culture media. Live-cell imaging recordings were conducted using a spinning disk microscope with incubating conditions of 37 °C with 5 % CO2. Cells were imaged every 2 sec using a 40 x oil objective. The raw image files were stored as 16-bit in “.nd2” format at 337x337 µm (1024x1024 pixel) at an interval of 2 sec and a total of 200 images per series (total time per series: 6:40 min). The image pre-processing was carried out with Fiji (ImageJ 1.52h) adapted from a previous publication.^99^ To compensate for photobleaching during the time series, the “Bleach Correction” tool was applied, followed by a top-hat spatial filter to increase gray values of mitochondrial objects. The total mitochondria count was calculated by creating a binary image where the minimum gray value (min) was set to min=Mean+StdDev and the maximum gray value (max) was set to max= Mean+StdDev+1, where “Mean” and “StdDev” were obtained by using the “Measure” tool. Next, the “watershed” tool was applied to break up large mitochondria networks. Total number of mitochondria was quantified using the “Analyze Particles” tool with standard settings and a size preference for 8-200 pixel. The moving mitochondria were defined by a particle size of at least 6 pixels that changed location over the time course of 4 frames (=8 seconds). This was accomplished by subtracting the following 4th frame for each frame in the time series (on the top-hat filtered image). Afterward, the subtracted image series was converted into a binary image series. The number of moving mitochondria was similarly counted using the “Analyze Particles” tool with standard settings and a size preference for 6-200 pixel. Finally, the percentage of moving mitochondria per sample was calculated by dividing the average number of moving mitochondria by the average number of total mitochondria, multiplied by 100. The percentage of stationary mitochondria was obtained by the subtraction of moving mitochondria from 100.

#### Western blotting

For western blots, 50-90 individual MO chronically treated with either DMSO (0.1 % v/v), 0.1 μM Sertaconazole or 1 μM Talarozole were collected on day 28-30 of differentiation in a 1.5 ml tube. MO were washed 3 times with PBS and pelleted using a tabletop centrifuge at 8,000 rpm for 5 min. Fresh pellets were weighed and resuspended in appropriate volume of RIPA Lysis Buffer supplemented with proteinase inhibitor and phosphatase inhibitor (1:100) cocktails. Samples were then transferred to tubes containing CK14 ceramic beads and processed using a tissue homogenizer. The suspension was incubated on ice for 30 min, followed by centrifugation for 10 min at 10,000 rcf at 4 °C. Protein levels in the supernatant were then quantified using the BCA Protein Assay Reagent Kit. Protein lysates (35 µg) were separated on 4–12 % Bis–Tris gels, transferred to PVDF membranes, which were blocked with 3 % skimmed milk powder in TBS-T and incubated with antibodies against SNPH, KIF5A and ACTB as a housekeeping protein (see **Key Resource table** for full list of antibodies). After overnight incubation at 4 °C, membranes were washed and incubated with secondary antibodies horseradish peroxidase-conjugated anti-rabbit or anti-mouse. Signals were visualized using ECL solutions and ChemiDoc Touch Imaging System and the intensity of the bands was quantified using Image Lab software (v. 6.1.0). The intensity of the bands of interest (SNPH and KIF5A) were normalized to ACTB content in the samples. Values of treated MO are expressed relative to DMSO-treated Leigh MO.

#### Targeted metabolomics

For targeted metabolomics analysis, NPCs were grown in Matrigel-coated 6-well plates until reaching 70 % confluence. NPCs were either left untreated or were treated with 0.1 % (v/v) DMSO, 10 μM Sertaconazole, or 10 μM Talarozole for 24 h. At this point, accutase was used to detach the cells from the plate. Cells were then centrifuged at 200 x *g* for 3 min and the obtained pellets were dissolved in PBS to allow for quantification of cells using an automatic cell counter. Using a gating size of 5-19 μm, 1 million NPCs/sample were pelleted in a 1.5 ml tube using a tabletop centrifuge at 8,000 rpm for 5 min. The supernatant was discarded, and the pellet was snap-frozen in liquid nitrogen for subsequent measurement. Extraction of the targeted metabolites was performed by adding 300 µl Methanol/H2O (80/20; v/v) to the 1 x 10^6^ cells, which were homogenized by ceramic beads CK14 using Precellys lysing kit. The corresponding isotopically labelled internal standards were added to the extraction solution. The targeted compounds were analyzed by UPLC-MS/MS. The system consists of a UPLC I-Class (Waters) coupled to a tandem mass spectrometer Xevo-TQ-XS (Waters). Electrospray ionization was performed in the negative ionization mode for the compounds fructose-1,6-bishphosphate, phosphoenolpyruvate (PEP) and 2/3-phosphoglycerate.^100^ The TCA, energy metabolites (AMP, ADP and ATP) as well acetyl- and succinyl-CoA were measured in the positive ionization mode.^101-103^ Mass spectrometric quantitation of the compounds was carried out in the multiple reaction monitoring (MRM) mode. MassLynx software (v. 4.2) was used for instrument’s control and data acquisition. Quantitation analysis was performed by TagetLynx XS software.

#### Cholesterol measurements

For cholesterol analysis, 40,000 NPCs per well were seeded onto a matrigel-coated 96-well black ibidi plates. The next day, NPCs were either fed with normal media (untreated condition) or treated with either 0.1 % (v/v) DMSO, 10 μM Sertaconazole, or 10 μM Talarozole for 48 h. For the 24 h treatment, NPCs were treated with the above-mentioned conditions two days after. Both 24 h and 48 h treatment conditions were fixed on the same day using 4% PFA for 15 min at RT and washed 3 times for 10 min with PBS. For PFO-GST labelling of membrane-bound cholesterol, fixed NPCs were blocked and permeabilized with 10 % goat serum in 0.1% Triton X-100, PBS (0.1 % PBST) for 1h and incubated with 15 µg/ml recombinant PFO-GST for 3 h.^104^ After washing the cells three times with 0.1 % PBST, anti-GST (1:200) was applied in 10% goat serum 0.1% PBST, overnight at 4 °C. After three washes with 0.1 % PBST, Alexa Fluor^TM^ 488 goat anti-mouse IgG (H+L) secondary antibody was applied (1:500) for 1 h at RT. Lastly, wells were washed three times in PBS and cells were preserved in Mounting Medium with DAPI. For neutral lipid labelling, fixed cells were incubated with 1 µg/mL BODIPY^TM^ 493/503 in PBS for 20 min, followed by three washes of 5 min with PBS and mounting medium with DAPI was added to the wells. Fluorescent images were acquired with a LSM 900 Zeiss confocal microscope. Laser power, gain and offset parameters were kept constant for each experiment, any subsequent adjustments to contrast and brilliance were applied equally to all images. The image analysis was performed using Fiji ImageJ software using custom-made macros.

#### Molecular docking

To test Sertaconazole and Talarozole binding to postulated targets, we performed *in silico* docking experiments. We modeled docking to the protein target of Talarozole cytochrome P45026 (CYP26A1/B1). Since there is no experimental structure reported to date for CYP26A1/B1, we obtained CYP26A1/B1 from the alphafold2^105^ database based on PDB:2VE3 which reports the structure for retinoic acid bound to cyanobacterial CYP120A1. Overlap between the two protein structures enabled the localization of the iron-heme and the substrate structure within the cavity of CYP26A1/B1.^106^ Removing the substrate and redocking it together with Sertaconazole and Talarozole revealed difference in poses and docking scores. CCDC Gold v2022 was used for docking and CCG MOE (v. 2022.02) was used to visualize the results and generate images. Compound poses were analyzed using docking scores and by the minimal distance of any compound’s atom from the heme-iron reactive center for oxidation. The combination of distance from the iron atom and the relative energy needed to reach the ligand’s position can suggest whether compounds act as substrate or as inhibitor, since inhibitors are usually closer to the iron-center disturbing the oxygen recruitment from heme.^107,108^

#### PCR and gene expression analyses

For quantitative real-time PCR (qPCR) analysis of CYP26B1 in iNs, 2 million NPCs were seeded onto matrigel-coated 6 well-plates one day prior to NGN2 lentiviral transfection supplemented with 4 μg/ml polybrene. After overnight incubation with NGN2 lentiviruses, cells were washed 3 times with PBS and the media was refreshed with sm+ containing 2 μg/ml of doxycycline to start neuronal induction (day 0). Three days later, media was exchanged to iN media (see *Neuromorphogenesis analysis in iNs* for media formulation) with either DMSO (0.1 % v/v), 10 μM Sertaconazole, or 10 μM Talarozole. On day 5, treatment was refreshed to incubate for another 6 h to observe changes in gene expression.^109^ Afterwards, cells were collected using a cell scraper, pelleted using a tabletop centrifuge at 8,000 rpm for 5 min, and snap-frozen in liquid nitrogen. RNA extraction from pellets was performed with RNeasy kit. cDNA was generated from 2 μg RNA using the first strand cDNA synthesis kit. qPCR experiments were conducted for three technical replicates and three biological iNs replicates with the SYBR green Mastermix and CFX96 Real-Time System (Bio-Rad). After averaging the CT values of the three technical replicates, the 2-ΔΔCT method was used. The expression levels of the genes of interest were normalized relative to the average expression of housekeeping genes (C1orf43 and YWHAZ) and relative to DMSO treatment. Primer sequences are reported in the **Key Resource Table**.^110^ Expression of anchoring and RA pathway genes was obtained from our previously published bulk RNAseq datasets for Leigh mDANs and Leigh CO.^14^

### QUANTIFICATION AND STATISTICAL ANALYSIS

Data are expressed as mean and standard deviation (mean ± SD) where normality of the distribution could be verified, or as median and quartiles (median [1^st^;4^th^ quartiles]) otherwise. Significance was assessed using parametric tests (Welch’s t-test, ANOVA) for normally distributed data and non-parametric tests (Mann-Whitney U test, Kruskal-Wallis) when normal distribution could not be verified. Unless otherwise indicated, data were analyzed and shown using GraphPad-Prism software (v. 10.2.2) (GraphPad Software, USA) or R. Schematic cartoons were made using Biorender.

---

**Figure S1. Neuromorphogenesis defects in 2D and 3D models of Leigh (related to Figure 1). (A-B)** Neurite outgrowth defects in Leigh iNs after NGN2 induction based on GFP signal that is part of the NGN2 inducible cassette. Mean + SD, dots represent 12 biological replicates over 2 independent experiments. Scale bars: 100 µm. **(C)** Schematic of midbrain organoid (MO) protocol. **(D)** Day 30 MO contained TH and NURR1-positive midbrain dopaminergic neurons (mDANs). Scale bars: 100 µm. **(E-K)** Functional calcium imaging in Leigh MO and CTL MO. Dots represent individual cells in MO (7 CTL MO and 6 Leigh MO) over 2 independent experiments; **p<0.01, ***p<0.005, ****p<0.001, Mann-Whitney U test.

**Figure S2. Neuromorphogenesis screenings in Leigh models with DL-predicted drugs (DLDs) and yeast screen drugs (YSDs) (related to Figure 2). (A)** UMAP showing unbiased clustering of CO scRNAseq. **(B)** Percent distribution of radial glia (RG) (cluster 7), intermediate progenitor cells (IPCs) (cluster 4) and neurons (cluster 6) at different pseudotime states in CO from Leigh and CTL. **(C)** Schematic of DL strategy. **(D-E)** Neuromorphogenesis screen in Leigh iNs based on HCA quantification of neurite outgrowth. Mean + SD of 5 biological replicates (dots); **p<0.01, ***p<0.005, ****p<0.001, unpaired two-tailed t test, compound-treated Leigh iNs vs DMSO-treated Leigh iNs. **(F-H)** Viability assay in cultures from wild-type (WT) yeast and from yeast knock-out for the SURF1 homologue SHY1 (ΔSHY). **(I)** FDA-approved drugs screened in ΔSHY yeasts. Dots represent individual drug response; green dots: top 50 yeast screen drugs (YSDs); purple dots: YSD selected for validations in Leigh iNs. **(J-K)** Neuromorphogenesis screen in Leigh iNs based on HCA quantification of neurite outgrowth. Mean + SD of 5 biological replicates (dots); **p<0.01, ***p<0.005, ****p<0.001, unpaired two-tailed t test, compound-treated Leigh iNs vs DMSO-treated Leigh iNs.

**Figure S3. Energy metabolism and mitochondrial anchoring in Leigh models (related to Figure 3 and Figure 4). (A-B)** Viability experiments in CTL MO showed that 0.1 µM Sertaconazole and 1 µM Talarzole were the highest concentrations that could be used for long-term treatment of MO. Dots represent individual MO over 3 independent experiments; ****p<0.001, unpaired two-tailed t test. **(C)** Representative images used to establish the pipeline to quantify moving and stationary mitochondria within midbrain dopaminergic neurons (mDANs) based on live cell tracking with MitoTracker Red. **(D)** Mitochondrial motility in CTL mDANs and Leigh mDANs upon acute exposure to the mitochondrial uncoupler FCCP. Dots represent individual mDANs over 3 independent experiments; ***p<0.005; unpaired two-tailed t test. **(E)** Left: Schematic depicting the relevance of anchoring proteins for mitochondrial motility. Right: expression of anchoring genes in Leigh mDANs and Leigh CO compared to CTL. *p<0.01, fold change expression from bulk RNAseq dataset. **(F-G)** tSNE plot of scRNAseq datasets from Leigh CO and CTL CO used for functional metabolic analysis showing different distribution of population with respect to all genes but overlapping profile for metabolism-related genes. **(H-K)** Functional metabolic changes in Leigh CO compared to CTL CO. Dots represent individual cell values for 3 independent CO scRNAseq samples.

**Figure S4. Mechanism of action studies in Leigh models treated with Sertaconazole or Talarozole (related to Figure 4). (A-D)** Targeted metabolomics for ATP and glycolysis-related metabolites in CTL vs Leigh NPCs and DMSO-treated Leigh NPCs vs. compound-treated Leigh NPCs. Dots represent biological replicates over 2 independent experiments; ns: not significant, **p<0.01, ***p<0.005, ****p<0.001; unpaired two-tailed t test. **(E-F)** Targeted metabolomics for TCA cycle related metabolites in CTL NPCs vs Leigh NPCs and DMSO-treated Leigh NPCs vs. compound-treated Leigh NPCs. Dots represent biological replicates over 2 independent experiments; ns: not significant, **p<0.01, ***p<0.005, ****p<0.001; unpaired two-tailed t test. **(G-H)** Quantification of neutral lipids stained with BODIPY in CTL NPCs and Leigh NPCs treated with Sertaconazole or Talarozole. Dots represent individual NPC samples over 3 independent experiments; ns not significant, ***p<0.005; unpaired two-tailed t test. Scale bars: 100 µm. **(I-K)** Docking modeling prediction for Sertaconazole and Talarozole with flexible configuration or with configuration fixed on the CYP120 pocket.

## References

1. Vafai, S.B., and Mootha, V.K. (2012). Mitochondrial disorders as windows into an ancient organelle. Nature 491, 374–383. 10.1038/nature11707.

2. Rahman, S. (2023). Leigh syndrome. Handb Clin Neurol 194, 43–63. 10.1016/B978-0-12-821751-1.00015-4.

3. Leigh, D. (1951). Subacute necrotizing encephalomyelopathy in an infant. J Neurol Neurosurg Psychiatry 14, 216–221.

4. Rahman, S., Blok, R.B., Dahl, H.H., Danks, D.M., Kirby, D.M., Chow, C.W., Christodoulou, J., and Thorburn, D.R. (1996). Leigh syndrome: clinical features and biochemical and DNA abnormalities. Ann Neurol 39, 343–351. 10.1002/ana.410390311.

5. Lake, N.J., Bird, M.J., Isohanni, P., and Paetau, A. (2015). Leigh syndrome: neuropathology and pathogenesis. J Neuropathol Exp Neurol 74, 482–492. 10.1097/NEN.0000000000000195.

6. Stenton, S.L., Zou, Y., Cheng, H., Liu, Z., Wang, J., Shen, D., Jin, H., Ding, C., Tang, X., Sun, S., et al. (2022). Leigh Syndrome: A Study of 209 Patients at the Beijing Children’s Hospital. Ann Neurol 91, 466–482. 10.1002/ana.26313.

7. 7. Zhu, Z., Yao, J., Johns, T., Fu, K., De Bie, I., Macmillan, C., Cuthbert, A.P., Newbold, R.F., Wang, J., Chevrette, M., et al. (1998). SURF1, encoding a factor involved in the biogenesis of cytochrome c oxidase, is mutated in Leigh syndrome. Nat Genet 20, 337–343. 10.1038/3804.

8. Wedatilake, Y., Brown, R.M., McFarland, R., Yaplito-Lee, J., Morris, A.A., Champion, M., Jardine, P.E., Clarke, A., Thorburn, D.R., Taylor, R.W., et al. (2013). SURF1 deficiency: a multi-centre natural history study. Orphanet J Rare Dis 8, 96. 10.1186/1750-1172-8-96.

9. Tyynismaa, H., and Suomalainen, A. (2009). Mouse models of mitochondrial DNA defects and their relevance for human disease. EMBO Rep 10, 137–143. 10.1038/embor.2008.242.

10. Dell’agnello, C., Leo, S., Agostino, A., Szabadkai, G., Tiveron, C., Zulian, A., Prelle, A., Roubertoux, P., Rizzuto, R., and Zeviani, M. (2007). Increased longevity and refractoriness to Ca(2+)-dependent neurodegeneration in Surf1 knockout mice. Hum Mol Genet 16, 431–444. 10.1093/hmg/ddl477.

11. Kovarova, N., Pecina, P., Nuskova, H., Vrbacky, M., Zeviani, M., Mracek, T., Viscomi, C., and Houstek, J. (2016). Tissue- and species-specific differences in cytochrome c oxidase assembly induced by SURF1 defects. Biochim Biophys Acta 1862, 705–715. 10.1016/j.bbadis.2016.01.007.

12. Russell, O.M., Gorman, G.S., Lightowlers, R.N., and Turnbull, D.M. (2020). Mitochondrial Diseases: Hope for the Future. Cell 181, 168–188. 10.1016/j.cell.2020.02.051.

13. Weissig, V. (2019). Drug Development for the Therapy of Mitochondrial Diseases. Trends Mol Med. 10.1016/j.molmed.2019.09.002.

14. Gerards, M., Sallevelt, S.C., and Smeets, H.J. (2016). Leigh syndrome: Resolving the clinical and genetic heterogeneity paves the way for treatment options. Mol Genet Metab 117, 300–312. 10.1016/j.ymgme.2015.12.004.

15. Dogan, S.A., Cerutti, R., Beninca, C., Brea-Calvo, G., Jacobs, H.T., Zeviani, M., Szibor, M., and Viscomi, C. (2018). Perturbed Redox Signaling Exacerbates a Mitochondrial Myopathy. Cell Metab 28, 764–775 e765. 10.1016/j.cmet.2018.07.012.

16. Inak, G., Rybak-Wolf, A., Lisowski, P., Pentimalli, T.M., Juttner, R., Glazar, P., Uppal, K., Bottani, E., Brunetti, D., Secker, C., et al. (2021). Defective metabolic programming impairs early neuronal morphogenesis in neural cultures and an organoid model of Leigh syndrome. Nat Commun 12, 1929. 10.1038/s41467-021-22117-z.

17. Brunetti, D., Dykstra, W., Le, S., Zink, A., and Prigione, A. (2021). Mitochondria in neurogenesis: Implications for mitochondrial diseases. Stem Cells. 10.1002/stem.3425.

18. Zhang, Y., Pak, C., Han, Y., Ahlenius, H., Zhang, Z., Chanda, S., Marro, S., Patzke, C., Acuna, C., Covy, J., et al. (2013). Rapid single-step induction of functional neurons from human pluripotent stem cells. Neuron 78, 785–798. 10.1016/j.neuron.2013.05.029.

19. Renner, H., Grabos, M., Becker, K.J., Kagermeier, T.E., Wu, J., Otto, M., Peischard, S., Zeuschner, D., TsyTsyura, Y., Disse, P., et al. (2020). A fully automated high-throughput workflow for 3D-based chemical screening in human midbrain organoids. Elife 9. 10.7554/eLife.52904.

20. Lickfett, S., Menacho, C., Zink, A., Telugu, N.S., Beller, M., Diecke, S., Cambridge, S., and Prigione, A. (2022). High-content analysis of neuronal morphology in human iPSC-derived neurons. STAR Protoc 3, 101567. 10.1016/j.xpro.2022.101567.

21. Zagare, A., Barmpa, K., Smajic, S., Smits, L.M., Grzyb, K., Grunewald, A., Skupin, A., Nickels, S.L., and Schwamborn, J.C. (2022). Midbrain organoids mimic early embryonic neurodevelopment and recapitulate LRRK2-p.Gly2019Ser-associated gene expression. Am J Hum Genet 109, 311–327. 10.1016/j.ajhg.2021.12.009.

22. Mashkevich, G., Repetto, B., Glerum, D.M., Jin, C., and Tzagoloff, A. (1997). SHY1, the yeast homolog of the mammalian SURF-1 gene, encodes a mitochondrial protein required for respiration. J Biol Chem 272, 14356–14364. 10.1074/jbc.272.22.14356.

23. Courchet, J., Lewis, T.L., Jr., Lee, S., Courchet, V., Liou, D.Y., Aizawa, S., and Polleux, F. (2013). Terminal axon branching is regulated by the LKB1-NUAK1 kinase pathway via presynaptic mitochondrial capture. Cell 153, 1510–1525. 10.1016/j.cell.2013.05.021.

24. Sheng, Z.H. (2014). Mitochondrial trafficking and anchoring in neurons: New insight and implications. J Cell Biol 204, 1087–1098. 10.1083/jcb.201312123.

25. Ilieva, M., Aldana, B.I., Vinten, K.T., Hohmann, S., Woofenden, T.W., Lukjanska, R., Waagepetersen, H.S., and Michel, T.M. (2022). Proteomic phenotype of cerebral organoids derived from autism spectrum disorder patients reveal disrupted energy metabolism, cellular components, and biological processes. Mol Psychiatry 27, 3749–3759. 10.1038/s41380-022-01627-2.

26 Pawel, L., Selene, L., Agnieszka, R.-W., Stephanie, L., Werner, D., barbara, m., Carmen, M., Philipp, R., Yasmin, R., Linda, K., et al. (2023). Mutant Huntingtin impairs neurodevelopment in human brain organoids through CHCHD2-mediated neurometabolic failure. bioRxiv, 2023.2006.2003.543551. 10.1101/2023.06.03.543551.

27. Fan, Q.W., Yu, W., Gong, J.S., Zou, K., Sawamura, N., Senda, T., Yanagisawa, K., and Michikawa, M. (2002). Cholesterol-dependent modulation of dendrite outgrowth and microtubule stability in cultured neurons. J Neurochem 80, 178–190. 10.1046/j.0022-3042.2001.00686.x.

28. Park, D.S., Kozaki, T., Tiwari, S.K., Moreira, M., Khalilnezhad, A., Torta, F., Olivie, N., Thiam, C.H., Liani, O., Silvin, A., et al. (2023). iPS-cell-derived microglia promote brain organoid maturation via cholesterol transfer. Nature 623, 397–405. 10.1038/s41586-023-06713-1.

29 Munoz-Oreja, M., Sandoval, A., Bruland, O., Perez-Rodriguez, D., Fernandez-Pelayo, U., de Arbina, A.L., Villar-Fernandez, M., Hernandez-Eguiazu, H., Hernandez, I., Park, Y., et al. (2024). Elevated cholesterol in ATAD3 mutants is a compensatory mechanism that leads to membrane cholesterol aggregation. Brain 147, 1899–1913. 10.1093/brain/awae018.

30. Croxtall, J.D., and Plosker, G.L. (2009). Sertaconazole: a review of its use in the management of superficial mycoses in dermatology and gynaecology. Drugs 69, 339–359. 10.2165/00003495-200969030-00009.

31. Ross, A.C., and Zolfaghari, R. (2011). Cytochrome P450s in the regulation of cellular retinoic acid metabolism. Annu Rev Nutr 31, 65–87. 10.1146/annurev-nutr-072610-145127.

32. Foti, R.S., Isoherranen, N., Zelter, A., Dickmann, L.J., Buttrick, B.R., Diaz, P., and Douguet, D. (2016). Identification of Tazarotenic Acid as the First Xenobiotic Substrate of Human Retinoic Acid Hydroxylase CYP26A1 and CYP26B1. J Pharmacol Exp Ther 357, 281–292. 10.1124/jpet.116.232637.

33. Kuhnel, K., Ke, N., Cryle, M.J., Sligar, S.G., Schuler, M.A., and Schlichting, I. (2008). Crystal structures of substrate-free and retinoic acid-bound cyanobacterial cytochrome P450 CYP120A1. Biochemistry 47, 6552–6559. 10.1021/bi800328s.

34. Tolle, I., Tiranti, V., and Prigione, A. (2023). Modeling mitochondrial DNA diseases: from base editing to pluripotent stem-cell-derived organoids. EMBO Rep 24, e55678. 10.15252/embr.202255678.

35. McKnight, C.L., Low, Y.C., Elliott, D.A., Thorburn, D.R., and Frazier, A.E. (2021). Modelling Mitochondrial Disease in Human Pluripotent Stem Cells: What Have We Learned? Int J Mol Sci 22. 10.3390/ijms22147730.

36. Bottani, E., Lamperti, C., Prigione, A., Tiranti, V., Persico, N., and Brunetti, D. (2020). Therapeutic Approaches to Treat Mitochondrial Diseases: “One-Size-Fits-All” and “Precision Medicine” Strategies. Pharmaceutics 12. 10.3390/pharmaceutics12111083.

37. Baertling, F., Rodenburg, R.J., Schaper, J., Smeitink, J.A., Koopman, W.J., Mayatepek, E., Morava, E., and Distelmaier, F. (2014). A guide to diagnosis and treatment of Leigh syndrome. J Neurol Neurosurg Psychiatry 85, 257–265. 10.1136/jnnp-2012-304426.

38. Chedotal, A., and Richards, L.J. (2010). Wiring the brain: the biology of neuronal guidance. Cold Spring Harb Perspect Biol 2, a001917. 10.1101/cshperspect.a001917.

39. Pacelli, C., Giguere, N., Bourque, M.J., Levesque, M., Slack, R.S., and Trudeau, L.E. (2015). Elevated Mitochondrial Bioenergetics and Axonal Arborization Size Are Key Contributors to the Vulnerability of Dopamine Neurons. Curr Biol 25, 2349–2360. 10.1016/j.cub.2015.07.050.

40. Bender, A., Schwarzkopf, R.M., McMillan, A., Krishnan, K.J., Rieder, G., Neumann, M., Elstner, M., Turnbull, D.M., and Klopstock, T. (2008). Dopaminergic midbrain neurons are the prime target for mitochondrial DNA deletions. J Neurol 255, 1231–1235 10.1007/s00415-008-0892-9.

41. Matsuda, W., Furuta, T., Nakamura, K.C., Hioki, H., Fujiyama, F., Arai, R., and Kaneko, T. (2009). Single nigrostriatal dopaminergic neurons form widely spread and highly dense axonal arborizations in the neostriatum. J Neurosci 29, 444–453. 10.1523/JNEUROSCI.4029-08.2009.

42. Lamb, J., Crawford, E.D., Peck, D., Modell, J.W., Blat, I.C., Wrobel, M.J., Lerner, J., Brunet, J.P., Subramanian, A., Ross, K.N., et al. (2006). The Connectivity Map: using gene-expression signatures to connect small molecules, genes, and disease. Science 313, 1929–1935. 10.1126/science.1132939.

43. Subramanian, A., Narayan, R., Corsello, S.M., Peck, D.D., Natoli, T.E., Lu, X., Gould, J., Davis, J.F., Tubelli, A.A., Asiedu, J.K., et al. (2017). A Next Generation Connectivity Map: L1000 Platform and the First 1,000,000 Profiles. Cell 171, 1437–1452 e1417. 10.1016/j.cell.2017.10.049.

44. Tong, X., Qu, N., Kong, X., Ni, S., Zhou, J., Wang, K., Zhang, L., Wen, Y., Shi, J., Zhang, S., et al. (2024). Deep representation learning of chemical-induced transcriptional profile for phenotype-based drug discovery. Nat Commun 15, 5378. 10.1038/s41467-024-49620-3.

45. Pawson, T., and Warner, N. (2007). Oncogenic re-wiring of cellular signaling pathways. Oncogene 26, 1268–1275. 10.1038/sj.onc.1210255.

46. Sharma, S., and Petsalaki, E. (2019). Large-scale datasets uncovering cell signalling networks in cancer: context matters. Curr Opin Genet Dev 54, 118–124. 10.1016/j.gde.2019.05.001.

47. Vander Heiden, M.G., Cantley, L.C., and Thompson, C.B. (2009). Understanding the Warburg effect: the metabolic requirements of cell proliferation. Science 324, 1029–1033. 10.1126/science.1160809.

48. Zheng, M., Okawa, S., Bravo, M., Chen, F., Martinez-Chantar, M.L., and Del Sol, A. (2023). ChemPert: mapping between chemical perturbation and transcriptional response for non-cancer cells. Nucleic Acids Res 51, D877–D889. 10.1093/nar/gkac862.

49. Tardiff, D.F., Jui, N.T., Khurana, V., Tambe, M.A., Thompson, M.L., Chung, C.Y., Kamadurai, H.B., Kim, H.T., Lancaster, A.K., Caldwell, K.A., et al. (2013). Yeast reveal a “druggable” Rsp5/Nedd4 network that ameliorates alpha-synuclein toxicity in neurons. Science 342, 979–983. 10.1126/science.1245321.

50 Couplan, E., Aiyar, R.S., Kucharczyk, R., Kabala, A., Ezkurdia, N., Gagneur, J., St Onge, R.P., Salin, B., Soubigou, F., Le Cann, M., et al. (2011). A yeast-based assay identifies drugs active against human mitochondrial disorders. Proc Natl Acad Sci U S A 108, 11989–11994. 10.1073/pnas.1101478108.

51. Chung, C.Y., Khurana, V., Auluck, P.K., Tardiff, D.F., Mazzulli, J.R., Soldner, F., Baru, V., Lou, Y., Freyzon, Y., Cho, S., et al. (2013). Identification and rescue of alpha-synuclein toxicity in Parkinson patient-derived neurons. Science 342, 983–987. 10.1126/science.1245296.

52. Njar, V.C., Gediya, L., Purushottamachar, P., Chopra, P., Vasaitis, T.S., Khandelwal, A., Mehta, J., Huynh, C., Belosay, A., and Patel, J. (2006). Retinoic acid metabolism blocking agents (RAMBAs) for treatment of cancer and dermatological diseases. Bioorg Med Chem 14, 4323–4340. 10.1016/j.bmc.2006.02.041.

53. Cunningham, T.J., and Duester, G. (2015). Mechanisms of retinoic acid signalling and its roles in organ and limb development. Nat Rev Mol Cell Biol 16, 110–123. 10.1038/nrm3932.

54. Gudas, L.J., and Wagner, J.A. (2011). Retinoids regulate stem cell differentiation. J Cell Physiol 226, 322–330. 10.1002/jcp.22417.

55. Diez del Corral, R., Olivera-Martinez, I., Goriely, A., Gale, E., Maden, M., and Storey, K. (2003). Opposing FGF and retinoid pathways control ventral neural pattern, neuronal differentiation, and segmentation during body axis extension. Neuron 40, 65–79. 10.1016/s0896-6273(03)00565-8.

56 Alekseenko, Z., Dias, J.M., Adler, A.F., Kozhevnikova, M., van Lunteren, J.A., Nolbrant, S., Jeggari, A., Vasylovska, S., Yoshitake, T., Kehr, J., et al. (2022). Robust derivation of transplantable dopamine neurons from human pluripotent stem cells by timed retinoic acid delivery. Nat Commun 13, 3046. 10.1038/s41467-022-30777-8.

57. Dmetrichuk, J.M., Carlone, R.L., and Spencer, G.E. (2006). Retinoic acid induces neurite outgrowth and growth cone turning in invertebrate neurons. Dev Biol 294, 39–49. 10.1016/j.ydbio.2006.02.018.

58. Clagett-Dame, M., McNeill, E.M., and Muley, P.D. (2006). Role of all-trans retinoic acid in neurite outgrowth and axonal elongation. J Neurobiol 66, 739–756. 10.1002/neu.20241.

59. Iwata, R., Casimir, P., Erkol, E., Boubakar, L., Planque, M., Gallego Lopez, I.M., Ditkowska, M., Gaspariunaite, V., Beckers, S., Remans, D., et al. (2023). Mitochondria metabolism sets the species-specific tempo of neuronal development. Science 379, eabn4705. 10.1126/science.abn4705.

60. Iwata, R., Casimir, P., and Vanderhaeghen, P. (2020). Mitochondrial dynamics in postmitotic cells regulate neurogenesis. Science 369, 858–862. 10.1126/science.aba9760.

61. Khacho, M., Harris, R., and Slack, R.S. (2019). Mitochondria as central regulators of neural stem cell fate and cognitive function. Nat Rev Neurosci 20, 34–48. 10.1038/s41583-018-0091-3.

62. Riancho, J., Berciano, M.T., Ruiz-Soto, M., Berciano, J., Landreth, G., and Lafarga, M. (2016). Retinoids and motor neuron disease: Potential role in amyotrophic lateral sclerosis. J Neurol Sci 360, 115–120. 10.1016/j.jns.2015.11.058.

63 Cunnane, S.C., Trushina, E., Morland, C., Prigione, A., Casadesus, G., Andrews, Z.B., Beal, M.F., Bergersen, L.H., Brinton, R.D., de la Monte, S., et al. (2020). Brain energy rescue: an emerging therapeutic concept for neurodegenerative disorders of ageing. Nat Rev Drug Discov 19, 609–633. 10.1038/s41573-020-0072-x.

64. Iqbal, L., Zameer, U., and Iqbal Malick, M. (2024). Exploring Talarozole as a Novel Therapeutic Approach for Osteoarthritis: Insights From Experimental Studies. Clin Med Insights Arthritis Musculoskelet Disord 17, 11795441231222494. 10.1177/11795441231222494.

65. Stevison, F., Jing, J., Tripathy, S., and Isoherranen, N. (2015). Role of Retinoic Acid-Metabolizing Cytochrome P450s, CYP26, in Inflammation and Cancer. Adv Pharmacol 74, 373–412. 10.1016/bs.apha.2015.04.006.

66. Zhu, L., Kamalathevan, P., Koneva, L.A., Zarebska, J.M., Chanalaris, A., Ismail, H., Wiberg, A., Ng, M., Muhammad, H., Walsby-Tickle, J., et al. (2022). Variants in ALDH1A2 reveal an anti-inflammatory role for retinoic acid and a new class of disease-modifying drugs in osteoarthritis. Sci Transl Med 14, eabm4054. 10.1126/scitranslmed.abm4054.

67. Garone, C., and Viscomi, C. (2018). Towards a therapy for mitochondrial disease: an update. Biochem Soc Trans 46, 1247–1261. 10.1042/BST20180134.

68. Steele, H., Gomez-Duran, A., Pyle, A., Hopton, S., Newman, J., Stefanetti, R.J., Charman, S.J., Parikh, J.D., He, L., Viscomi, C., et al. (2020). Metabolic effects of bezafibrate in mitochondrial disease. EMBO Mol Med 12, e11589. 10.15252/emmm.201911589.

69. Wang, X., Sterr, M., Burtscher, I., Chen, S., Hieronimus, A., Machicao, F., Staiger, H., Haring, H.U., Lederer, G., Meitinger, T., et al. (2018). Genome-wide analysis of PDX1 target genes in human pancreatic progenitors. Mol Metab 9, 57–68. 10.1016/j.molmet.2018.01.011.

70. Butler, A., Hoffman, P., Smibert, P., Papalexi, E., and Satija, R. (2018). Integrating single-cell transcriptomic data across different conditions, technologies, and species. Nat Biotechnol 36, 411–420. 10.1038/nbt.4096.

71. Qiu, X., Mao, Q., Tang, Y., Wang, L., Chawla, R., Pliner, H.A., and Trapnell, C. (2017). Reversed graph embedding resolves complex single-cell trajectories. Nat Methods 14, 979–982. 10.1038/nmeth.4402.

72. Langer, J., and Rose, C.R. (2009). Synaptically induced sodium signals in hippocampal astrocytes in situ. J Physiol 587, 5859–5877. 10.1113/jphysiol.2009.182279.

73. Razick, S., Magklaras, G., and Donaldson, I.M. (2008). iRefIndex: a consolidated protein interaction database with provenance. BMC Bioinformatics 9, 405. 10.1186/1471-2105-9-405.

74. Turei, D., Korcsmaros, T., and Saez-Rodriguez, J. (2016). OmniPath: guidelines and gateway for literature-curated signaling pathway resources. Nat Methods 13, 966–967. 10.1038/nmeth.4077.

75. Wu, G., Feng, X., and Stein, L. (2010). A human functional protein interaction network and its application to cancer data analysis. Genome Biol 11, R53. 10.1186/gb-2010-11-5-r53.

76. Kanehisa, M., and Goto, S. (2000). KEGG: kyoto encyclopedia of genes and genomes. Nucleic Acids Res 28, 27–30. 10.1093/nar/28.1.27.

77. Browaeys, R., Saelens, W., and Saeys, Y. (2020). NicheNet: modeling intercellular communication by linking ligands to target genes. Nat Methods 17, 159–162. 10.1038/s41592-019-0667-5.

78. Hou, R., Denisenko, E., Ong, H.T., Ramilowski, J.A., and Forrest, A.R.R. (2020). Predicting cell-to-cell communication networks using NATMI. Nat Commun 11, 5011. 10.1038/s41467-020-18873-z.

79. Ximerakis, M., Lipnick, S.L., Innes, B.T., Simmons, S.K., Adiconis, X., Dionne, D., Mayweather, B.A., Nguyen, L., Niziolek, Z., Ozek, C., et al. (2019). Single-cell transcriptomic profiling of the aging mouse brain. Nat Neurosci 22, 1696–1708. 10.1038/s41593-019-0491-3.

80. Ramilowski, J.A., Goldberg, T., Harshbarger, J., Kloppmann, E., Lizio, M., Satagopam, V.P., Itoh, M., Kawaji, H., Carninci, P., Rost, B., and Forrest, A.R. (2015). A draft network of ligand-receptor-mediated multicellular signalling in human. Nat Commun 6, 7866. 10.1038/ncomms8866.

81. Shao, X., Liao, J., Li, C., Lu, X., Cheng, J., and Fan, X. (2021). CellTalkDB: a manually curated database of ligand-receptor interactions in humans and mice. Brief Bioinform 22. 10.1093/bib/bbaa269.

82. Wang, J., Wang, J., Hu, M., Wu, S., Qi, J., Wang, G., Han, Z., Qi, Y., Gao, N., Wang, H.W., et al. (2019). Ligand-triggered allosteric ADP release primes a plant NLR complex. Science 364. 10.1126/science.aav5868.

83. Jin, S., Guerrero-Juarez, C.F., Zhang, L., Chang, I., Ramos, R., Kuan, C.H., Myung, P., Plikus, M.V., and Nie, Q. (2021). Inference and analysis of cell-cell communication using CellChat. Nat Commun 12, 1088. 10.1038/s41467-021-21246-9.

84. Cabello-Aguilar, S., Alame, M., Kon-Sun-Tack, F., Fau, C., Lacroix, M., and Colinge, J. (2020). SingleCellSignalR: inference of intercellular networks from single-cell transcriptomics. Nucleic Acids Res 48, e55. 10.1093/nar/gkaa183.

85. Qiao, W., Wang, W., Laurenti, E., Turinsky, A.L., Wodak, S.J., Bader, G.D., Dick, J.E., and Zandstra, P.W. (2014). Intercellular network structure and regulatory motifs in the human hematopoietic system. Mol Syst Biol 10, 741. 10.15252/msb.20145141.

86. Vento-Tormo, R., Efremova, M., Botting, R.A., Turco, M.Y., Vento-Tormo, M., Meyer, K.B., Park, J.E., Stephenson, E., Polanski, K., Goncalves, A., et al. (2018). Single-cell reconstruction of the early maternal-fetal interface in humans. Nature 563, 347–353. 10.1038/s41586-018-0698-6.

87. Noel, F., Massenet-Regad, L., Carmi-Levy, I., Cappuccio, A., Grandclaudon, M., Trichot, C., Kieffer, Y., Mechta-Grigoriou, F., and Soumelis, V. (2021). Dissection of intercellular communication using the transcriptome-based framework ICELLNET. Nat Commun 12, 1089. 10.1038/s41467-021-21244-x.

88. Pavlicev, M., Wagner, G.P., Chavan, A.R., Owens, K., Maziarz, J., Dunn-Fletcher, C., Kallapur, S.G., Muglia, L., and Jones, H. (2017). Single-cell transcriptomics of the human placenta: inferring the cell communication network of the maternal-fetal interface. Genome Res 27, 349–361. 10.1101/gr.207597.116.

89. Hu, H., Miao, Y.R., Jia, L.H., Yu, Q.Y., Zhang, Q., and Guo, A.Y. (2019). AnimalTFDB 3.0: a comprehensive resource for annotation and prediction of animal transcription factors. Nucleic Acids Res 47, D33–D38. 10.1093/nar/gky822.

90. Dull, T., Zufferey, R., Kelly, M., Mandel, R.J., Nguyen, M., Trono, D., and Naldini, L. (1998). A third-generation lentivirus vector with a conditional packaging system. J Virol 72, 8463–8471. 10.1128/JVI.72.11.8463-8471.1998.

91. Busskamp, V., Lewis, N.E., Guye, P., Ng, A.H., Shipman, S.L., Byrne, S.M., Sanjana, N.E., Murn, J., Li, Y., Li, S., et al. (2014). Rapid neurogenesis through transcriptional activation in human stem cells. Mol Syst Biol 10, 760. 10.15252/msb.20145508.

92. Vierbuchen, T., Ostermeier, A., Pang, Z.P., Kokubu, Y., Sudhof, T.C., and Wernig, M. (2010). Direct conversion of fibroblasts to functional neurons by defined factors. Nature 463, 1035–1041. 10.1038/nature08797.

93. Carpenter, A.E., Jones, T.R., Lamprecht, M.R., Clarke, C., Kang, I.H., Friman, O., Guertin, D.A., Chang, J.H., Lindquist, R.A., Moffat, J., et al. (2006). CellProfiler: image analysis software for identifying and quantifying cell phenotypes. Genome Biol 7, R100. 10.1186/gb-2006-7-10-r100.

94. Berndt, N., Eckstein, J., Wallach, I., Nordmeyer, S., Kelm, M., Kirchner, M., Goubergrits, L., Schafstedde, M., Hennemuth, A., Kraus, M., et al. (2021). CARDIOKIN1: Computational Assessment of Myocardial Metabolic Capability in Healthy Controls and Patients With Valve Diseases. Circulation 144, 1926–1939. 10.1161/CIRCULATIONAHA.121.055646.

95. Berndt, N., Kann, O., and Holzhutter, H.G. (2015). Physiology-based kinetic modeling of neuronal energy metabolism unravels the molecular basis of NAD(P)H fluorescence transients. J Cereb Blood Flow Metab 35, 1494–1506. 10.1038/jcbfm.2015.70.

96. Bulik, S., Holzhutter, H.G., and Berndt, N. (2016). The relative importance of kinetic mechanisms and variable enzyme abundances for the regulation of hepatic glucose metabolism--insights from mathematical modeling. BMC Biol 14, 15. 10.1186/s12915-016-0237-6.

97. Berndt, N., Bulik, S., Wallach, I., Wunsch, T., Konig, M., Stockmann, M., Meierhofer, D., and Holzhutter, H.G. (2018). HEPATOKIN1 is a biochemistry-based model of liver metabolism for applications in medicine and pharmacology. Nat Commun 9, 2386. 10.1038/s41467-018-04720-9.

98. Reinhardt, P., Glatza, M., Hemmer, K., Tsytsyura, Y., Thiel, C.S., Hoing, S., Moritz, S., Parga, J.A., Wagner, L., Bruder, J.M., et al. (2013). Derivation and expansion using only small molecules of human neural progenitors for neurodegenerative disease modeling. PLoS One 8, e59252. 10.1371/journal.pone.0059252.

99. Iannetti, E.F., Smeitink, J.A., Beyrath, J., Willems, P.H., and Koopman, W.J. (2016). Multiplexed high-content analysis of mitochondrial morphofunction using live-cell microscopy. Nat Protoc 11, 1693–1710. 10.1038/nprot.2016.094.

100 Campos, C.G., Veras, H.C.T., de Aquino Ribeiro, J.A., Costa, P., Araujo, K.P., Rodrigues, C.M., de Almeida, J.R.M., and Abdelnur, P.V. (2017). New Protocol Based on UHPLC-MS/MS for Quantitation of Metabolites in Xylose-Fermenting Yeasts. J Am Soc Mass Spectrom 28, 2646–2657. 10.1007/s13361-017-1786-9.

101. Marquis, B.J., Louks, H.P., Bose, C., Wolfe, R.R., and Singh, S.P. (2017). A New Derivatization Reagent for HPLC-MS Analysis of Biological Organic Acids. Chromatographia 80, 1723–1732. 10.1007/s10337-017-3421-0.

102. Law, A.S., Hafen, P.S., and Brault, J.J. (2022). Liquid chromatography method for simultaneous quantification of ATP and its degradation products compatible with both UV-Vis and mass spectrometry. J Chromatogr B Analyt Technol Biomed Life Sci 1206, 123351. 10.1016/j.jchromb.2022.123351.

103. Jones, A.E., Arias, N.J., Acevedo, A., Reddy, S.T., Divakaruni, A.S., and Meriwether, D. (2021). A Single LC-MS/MS Analysis to Quantify CoA Biosynthetic Intermediates and Short-Chain Acyl CoAs. Metabolites 11. 10.3390/metabo11080468.

104. Goicoechea, L., Arenas, F., Castro, F., Nunez, S., Torres, S., Garcia-Ruiz, C., and Fernandez-Checa, J.C. (2022). GST-Perfringolysin O production for the localization and quantification of membrane cholesterol in human and mouse brain and liver. STAR Protoc 3, 101068. 10.1016/j.xpro.2021.101068.

105. Jumper, J., Evans, R., Pritzel, A., Green, T., Figurnov, M., Ronneberger, O., Tunyasuvunakool, K., Bates, R., Zidek, A., Potapenko, A., et al. (2021). Highly accurate protein structure prediction with AlphaFold. Nature 596, 583–589. 10.1038/s41586-021-03819-2.

106. Jones, G., Willett, P., Glen, R.C., Leach, A.R., and Taylor, R. (1997). Development and validation of a genetic algorithm for flexible docking. J Mol Biol 267, 727–748. 10.1006/jmbi.1996.0897.

107 Famulari, A., Correddu, D., Di Nardo, G., Gilardi, G., Mitrikas, G., Chiesa, M., and Garcia-Rubio, I. (2024). Heme Spin Distribution in the Substrate-Free and Inhibited Novel CYP116B5hd: A Multifrequency Hyperfine Sublevel Correlation (HYSCORE) Study. Molecules 29. 10.3390/molecules29020518.

108. Xu, L., and Chen, L.Y. (2020). Molecular determinant of substrate binding and specificity of cytochrome P450 2J2. Sci Rep 10, 22267. 10.1038/s41598-020-79284-0.

109. Stevison, F., Hogarth, C., Tripathy, S., Kent, T., and Isoherranen, N. (2017). Inhibition of the all-trans Retinoic Acid (atRA) Hydroxylases CYP26A1 and CYP26B1 Results in Dynamic, Tissue-Specific Changes in Endogenous atRA Signaling. Drug Metab Dispos 45, 846–854. 10.1124/dmd.117.075341.

110. Artyukhov, A.S., Dashinimaev, E.B., Tsvetkov, V.O., Bolshakov, A.P., Konovalova, E.V., Kolbaev, S.N., Vorotelyak, E.A., and Vasiliev, A.V. (2017). New genes for accurate normalization of qRT-PCR results in study of iPS and iPS-derived cells. Gene 626, 234–240. 10.1016/j.gene.2017.05.045.

