## Supplementary figures and images for "Deep learning-driven neuromorphogenesis screenings identify repurposable drugs for mitochondrial disease"

### S1

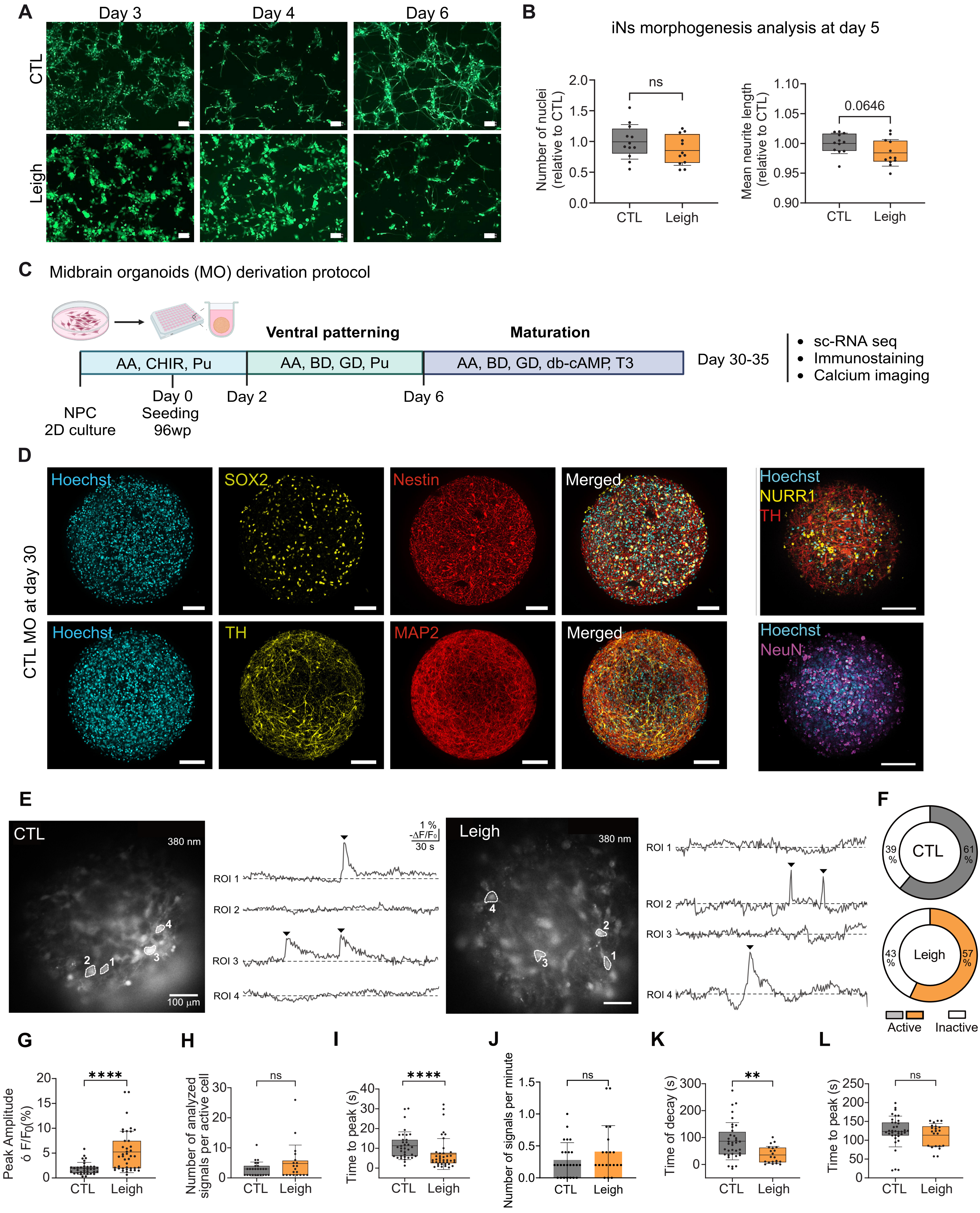

### S2

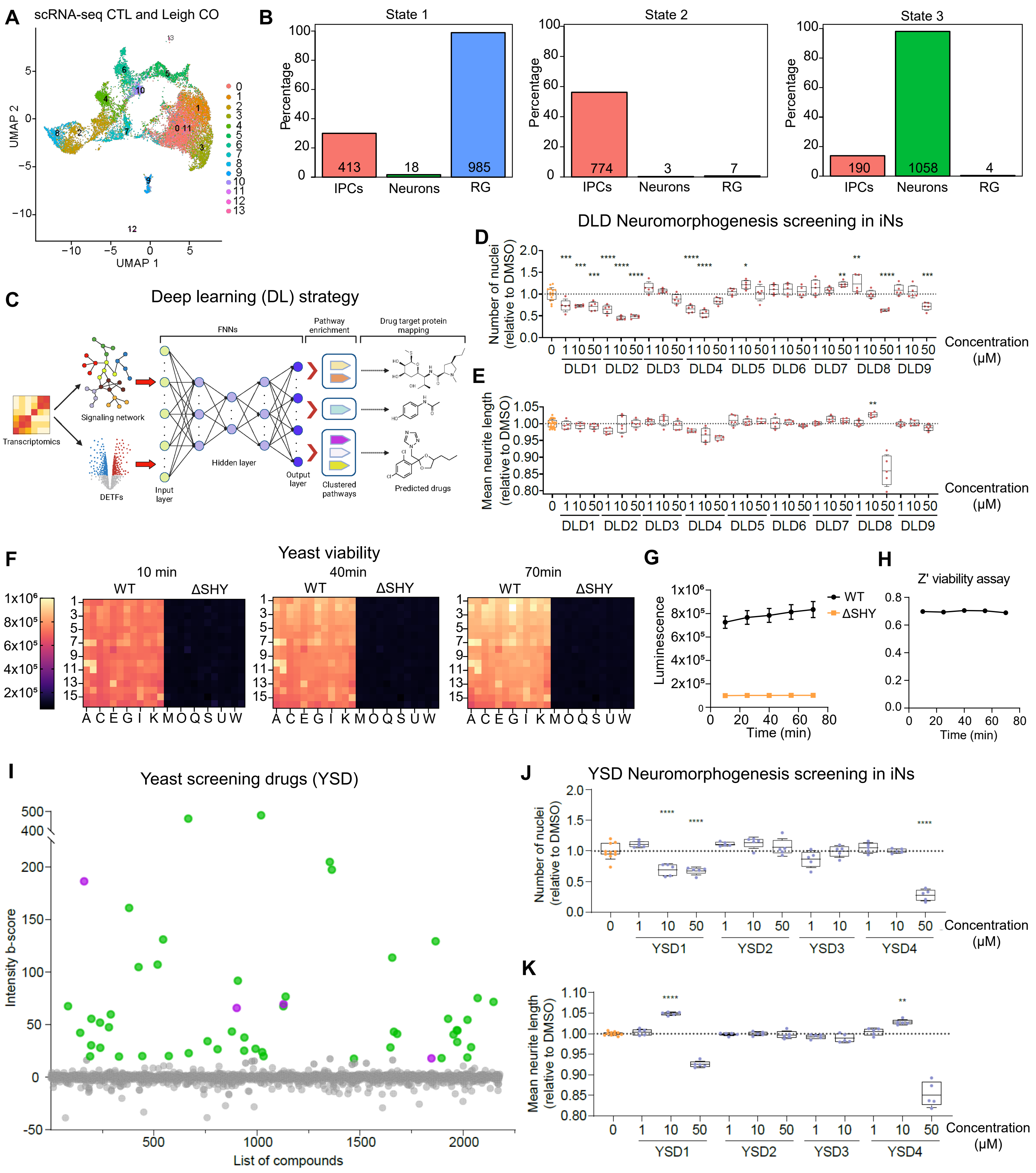

### S3

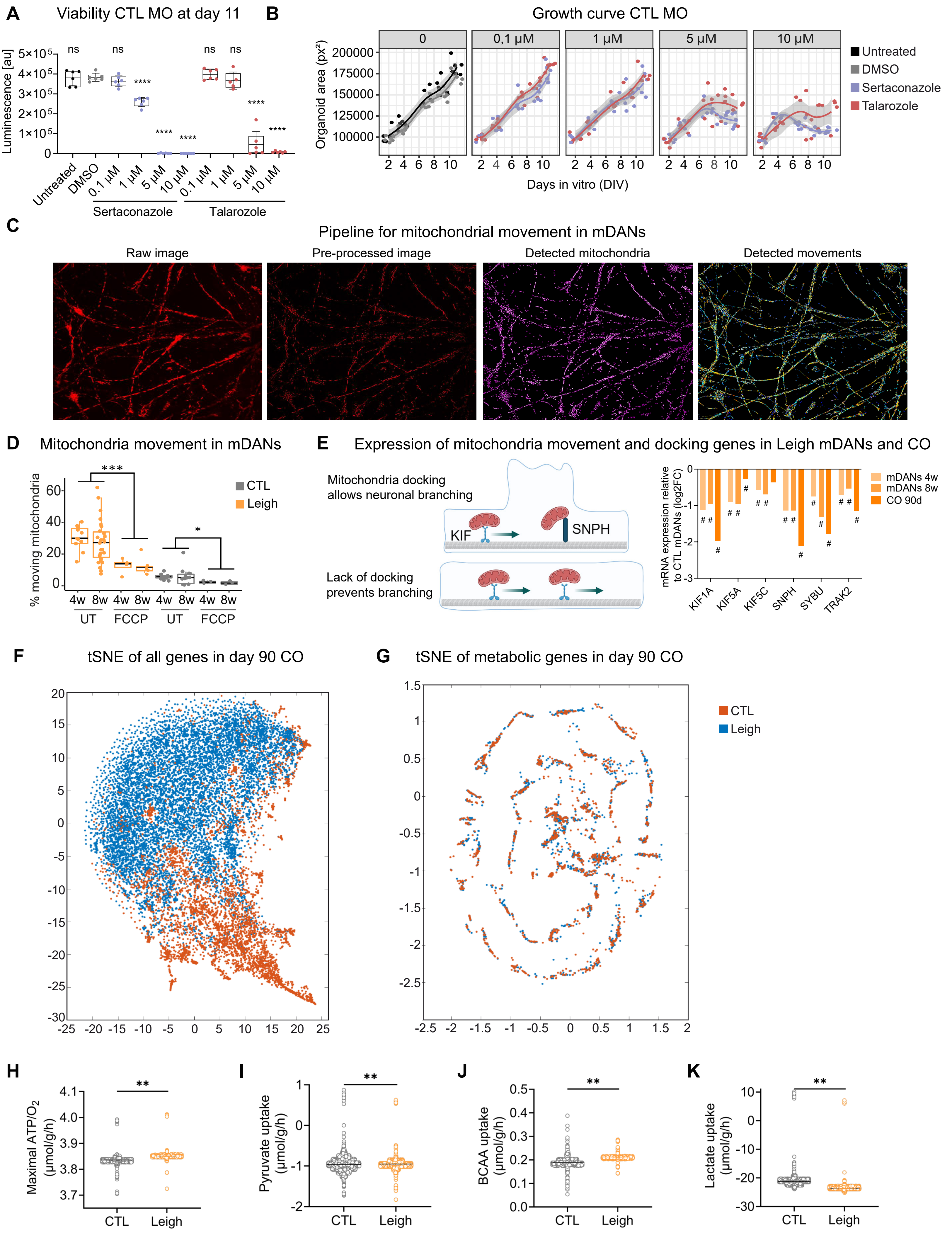

### S4

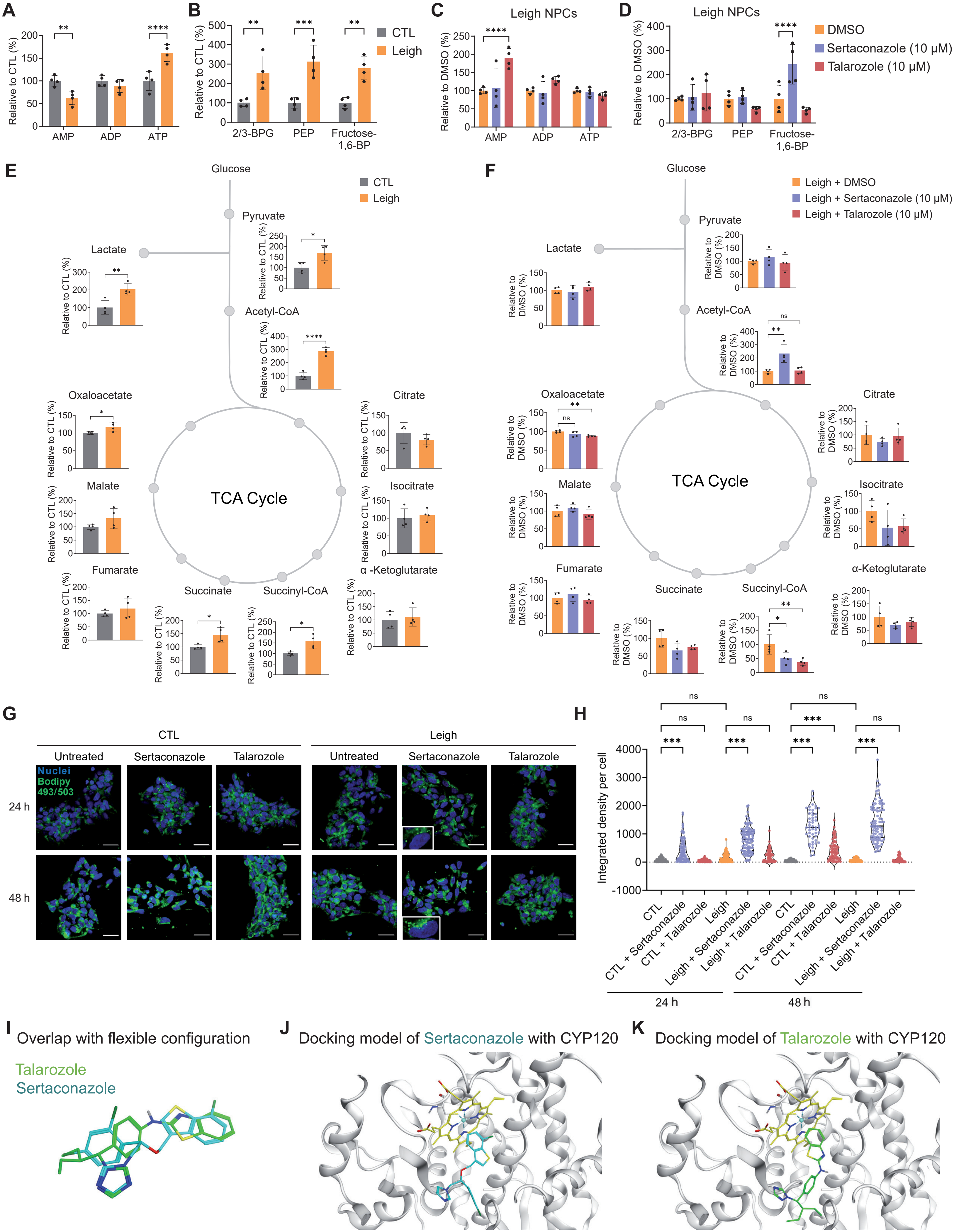
